# Structural Brain Indicators of Cognitive Performance in Middle and Late Adulthood: The Human Connectome Project in Aging/Aging Adult Brain Connectome Cohort

**DOI:** 10.64898/2026.01.21.700887

**Authors:** Yue Hong, Christa Michel, Gabriele Maria Gassner, Courtney Accorsi, Beau M Ances, Lauren M Antonucci, Steven E Arnold, Ganesh M Babulal, Susan Y Bookheimer, Randy L Buckner, Carlos Cruchaga, Mirella Díaz-Santos, Jennifer Stine Elam, David Van Essen, Dara G Ghahremani, Matthew F Glasser, Michael P Harms, Meher R Juttukonda, Adam M Khay, Jan A Kufer, Helen Lavretsky, Petra Lenzini, Ross W Mair, Pauline M Maki, Thomas E Nichols, Angela Oliver, Eva M Ratai, Gayathri Vijayaraghavan, Robert C Welsh, Essa Yacoub, Yiwen Zhang, David H Salat, Aging Adult Brain Connectome (AABC) Consortium

**Affiliations:** Athinoula A. Martinos Center for Biomedical Imaging, Massachusetts General Hospital, Boston, MA, USA; Department of Radiology, Harvard Medical School, Boston, MA, USA; Department of Neurology, Washington University School of Medicine, St Louis, MO, USA; Department of Neurology, Massachusetts General Hospital, Boston, MA 02114, USA; Department of Neurology, Harvard Medical School, Boston, MA 02114, USA; Institute of Public Health, Washington University in St. Louis, St. Louis, Missouri, USA; Department of Psychiatry and Biobehavioral Sciences, Semel Institute for Neuroscience and Human Behavior, David Geffen School of Medicine, University of California, Los Angeles, Los Angeles, CA, USA; Division of Medical Sciences, Harvard Medical School, Boston, Massachusetts, USA; Center for Brain Science, Harvard University, Cambridge, Massachusetts, USA; Department of Psychiatry, Massachusetts General Hospital, Charlestown, Massachusetts, USA; Department of Psychiatry, Washington University School of Medicine, St. Louis, MO, USA; NeuroGenomics and Informatics Center, Washington University, St. Louis, MO, USA; Mary S Easton Center for Alzheimer’s Research and Care, Department of Neurology, University of California, Los Angeles, Los Angeles, CA, USA; Department of Neuroscience, Washington University School of Medicine, St. Louis, MO, USA; Department of Radiology, Washington University School of Medicine, St. Louis, MO, USA; Departments of Psychiatry, Psychology, and OB/GYN, University of Illinois at Chicago, Chicago, Illinois, USA; Big Data Institute, Li Ka Shing Centre for Health Information and Discovery, Nuffield Department of Medicine, University of Oxford, Oxford, UK; Center for Magnetic Resonance Research (CMRR), University of Minnesota, Minneapolis, MN, USA; Carnegie Mellon University Neuroscience Institute, Pittsburgh, PA, USA; Centre for Integrative Neuroimaging, FMRIB, Nuffield Department of Clinical Neurosciences, University of Oxford, Oxford, OX3 9DU, UK; Department of Neurology, University Hospital Zurich, Zurich, Switzerland

## Abstract

**Introduction:** Significant effort has been put towards mapping patterns of atrophy in the cerebral cortex that are related to pathological aging, including the characteristic patterns of neurodegeneration in Alzheimer’s disease (AD). In contrast, brain structural patterns that support preserved or even exceptional cognition throughout the adult age-span and especially in later life are much less known. It is possible that superior cognitive performance in late life is supported by a preservation of brain structures vulnerable to typical aging. Alternatively, elevated performance could be related to preservation of brain regions vulnerable to age-associated pathology. Examination of individuals that exhibit superior cognition throughout the adult lifespan may provide unique insights into neural mechanisms that support cognitive resilience in late life.

**Methods:** We examined cross-sectional associations between cortical brain structure and cognitive performance across three stages of adulthood: midlife (36-59), young-old (60-79), and older adults (80+) in typically aging individuals enrolled as part of the Human Connectome Project Lifespan-Aging (HCP-A)/Aging Adult Brain Connectome (AABC) studies. Participants were considered generally healthy and excluded for significant and/or atypical health conditions for their demographic category, including a clinical diagnosis of cognitive impairment or dementia. Domain-specific cognitive factor scores representing memory, fluid intelligence, and crystalized intelligence were sex-stratified and residualized relative to age, and participants were classified as high, middle, or low performers based on their unique performance relative to the study sample.

**Results:** In the full sample, high performers demonstrated greater cortical thickness in regions of somatomotor, visual, and auditory cortices, as well as cortical areas in the frontal, parietal, and insular cortices (e.g., 5m, LIPv, MBelt). We also found associations between medial temporal and cingulate cortical thickness and cognitive performance, but only for select analyses. Group differences in cortical thickness were greatest when contrasting high and low performers for fluid intelligence compared to the other cognitive factors and were most prominent in the midlife participants compared to the other age strata. These group differences were primarily driven by reduced cortical thickness in the low performing individuals in the younger age bins relative to the ‘typically’ performing sample. Effects were rather limited when contrasting high and low performers in the 80+ age group. The cortical areas of high statistical significance appeared to show a cross-sectional convergence effect, such that the differences in cortical thickness between high vs. low cognitive performance groups diminished with increasing age. Despite lower statistical power, effect sizes were greatest when contrasting individuals at the extremes of performance (e.g., top 10% vs. bottom 10% performers). These effects were robust to subsample replications using longitudinally defined cognitive classifications.

**Discussion:** Elevated cognitive performance was cross-sectionally associated with increased regional cortical thickness and effects were most prominent in mid-life compared to later ages. Notably, contrary to brain regions that may be expected to support such high-order cognitive performance, a significant portion of primary sensory, motor, and insular cortical areas exhibited group differences between the high- and low-performing groups in this relatively younger age group. Group differences were due to lower thickness in these regions in the low performing group relative to the typical performers. In contrast to the younger portion of the sample, regions typically considered vulnerable to Alzheimer’s disease (i.e., regions in the medial temporal lobe) were only infrequently implicated. These results suggest that specific patterns of cortical brain structural integrity including preserved thickness of primary motor and sensory cortical regions may be a necessary but not sufficient mechanism supporting superior cognitive abilities in earlier adulthood, while alternate neural mechanisms may support cognitive resilience later in life. These results must be interpreted with caution given the cross-sectional nature of this study. Cognitive capacity can only be estimated from a single timepoint, and several factors contribute to inter-individual variation that will not be accounted for in the models applied here. Longitudinal assessment of cognitive resilience in the HCP-A/AABC cohort will be performed in future work.

## 1. Introduction

The prevalence of cognitively debilitating conditions resulting in impairment and dementia is rising as the average life expectancy increases ^1^. Significant effort has been put towards mapping patterns of atrophy in the cerebral cortex that are related to age-associated diseases such as the hallmark pathology of neurodegeneration in Alzheimer’s disease (AD) and related dementia (ADRD) ^2–10^. In contrast, less is known about mechanisms that promote preserved and even superior cognitive abilities in late life. Recent studies linking cortical structural integrity to successful aging or “super-aging” have emphasized the cingulate cortex^11–17^, and emerging reports also point to broader involvement of additional regions (e.g., insula and prefrontal cortex)^15,18–20^ and distributed networks (e.g., default mode and salience networks)^21,22^. Together, this literature has advanced the field substantially. At the same time, many studies have necessarily relied on modest samples for whole-brain analyses and on statistical approaches that can make it challenging to fully separate age effects from the intertwined relations among age, brain structure, and cognitive performance^23–25^. Several key questions remain. Does better-than-expected cognitive performance in late life reflect preservation of brain structures most vulnerable to ‘typical’ aging, including frontal, prefrontal, sensory, and motor areas ^23,26,27^? Or is superior cognitive functioning more closely tied to preserved neural integrity in regions that are selectively vulnerable to AD-related pathology, such as the hippocampus, parahippocampal regions and entorhinal cortex in the medial temporal lobe ^7–9,28–32^? It is also possible that the relationship between cognitive performance and neural integrity changes over the course of aging, such that mechanisms supporting cognition in younger adults are not the same as those in older adults^13,33^.

Examination of individuals in the latest decades of life that exhibit preserved, even superior, cognition may provide unique insights into mechanisms that support late-life cognitive resilience. Building on the body of work demonstrating that structural brain measures are associated with cognitive performance across the adult lifespan ^11,12,15,19,25,34–40^, we investigated associations between cortical thickness and cognitive performance in cognitively healthy individuals characterized as high or low performers across the mid-to-late adult lifespan, based on three proxies of cognitive domain functioning derived through factor analysis. We employed methods to carefully control the confounding effects of age. We use Imaging Derived Phenotype (IDP, i.e., region of interest)-based linear models in a large cross-sectional community sample of cognitively intact individuals from the Lifespan Human Connectome Project in Aging (HCP-A) and Aging Adult Brain Connectome (AABC) cohorts to determine 1) whether continuous variation in cortical thickness is associated with cognitive performance; 2) whether individuals at the extremes of cognitive performance (high and low performers) show unique associations with cortical thickness compared to the overall continuous variation, and 3) whether relationships between cortical thickness and cognitive performance differ across age-strata in mid-to-late adult lifespan. We hypothesized that increasing cognitive ability would be associated with increasing cortical thickness, and that top cognitive performers would exhibit regionally thicker cortices compared to bottom performers.

## 2. Methods

### 2.1 Human Connectome Project – Aging: Participant Data

The Lifespan Human Connectome Project in Aging/Aging Adult Brain Connectome (HCP-A/AABC) study followed adults aged 36 – 90+ who were cognitively healthy at baseline across four acquisition sites using matched neuroimaging and characterization protocols^41,42^. The AABC is the ongoing second phase of the HCP-A project that includes additional study visits of the HCP-A cohort using protocols primarily matched to the HCP-A protocols with some modifications including acquisition of additional data types, as well as enrollment of a new cohort of baseline participants. All volunteers provided informed consent and Institutional Review Board approval was obtained.

Individuals with a diagnosis of stroke, dementia or cognitive impairment, or any major neurological or psychological disorder were excluded from participation. The Montreal Cognitive Assessment (MoCA) was used to exclude individuals with gross cognitive impairment using a minimum required score that varied by age range (19 for individuals aged 36 – <80, 17 for individuals aged 80 – <90, and 16 for individuals aged 90+). Full details on the study protocol have been described previously^41,42^. The present study used a cross-sectional sample of HCP-A/AABC data collected prior to November 2023, with an internal data release date of April 2024. The data release sample described here includes only baseline visits of participants who have a complete set of cognitive data and whose T1-weighted images passed quality control (see Table 1).

**Table 1.**
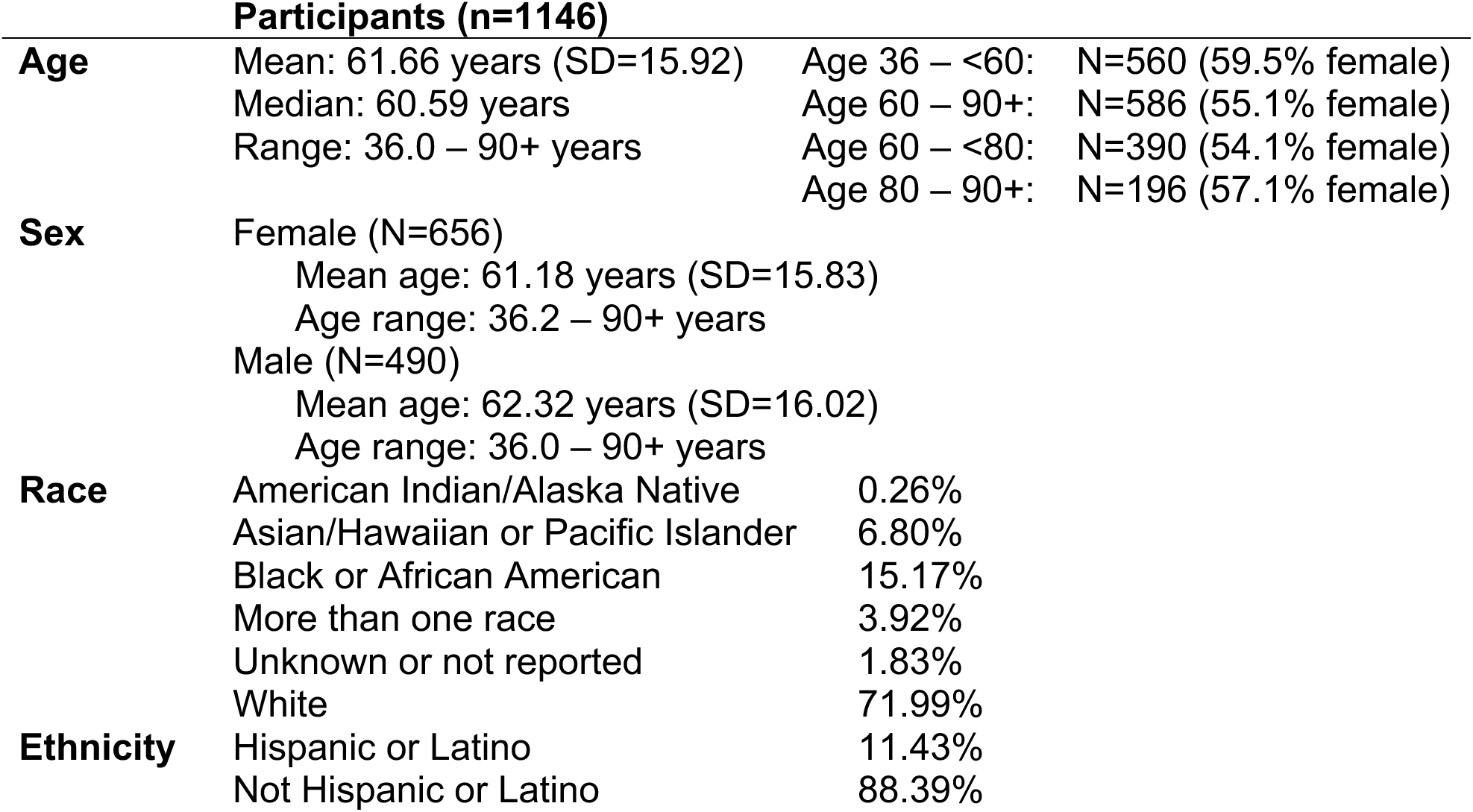
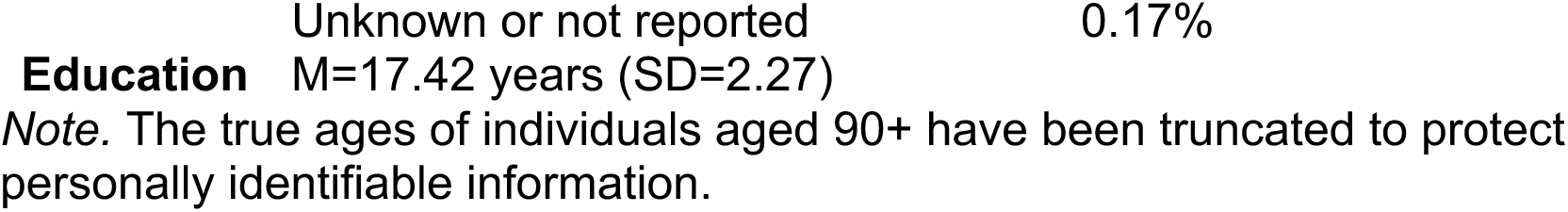
Sample demographics.

### 2.2 Neuropsychological Data

The complete neuropsychological assessment for HCP-A/AABC data has been described previously^41^. HCP-A/AABC participants aged between 36 and 80 years old with complete neuropsychological data (N=962, 552 female; mean age = 56.7 years, SD = 12.44 years) were included in a factor analysis to reduce the battery to primary cognitive dimensions. These loadings were then used to compute factor scores for participants above the age of 80. The factor analysis was performed on 25 variables from the neuropsychological battery. Minimum residual factor extraction was used to extract 3 dimensions that maximally captured non-random structure in the data and appeared to capture different domains of cognitive ability (Figure 1). The full list of neuropsychological tests included in the factor analysis and each factor loading is available in Table S1.

**Figure 1.**
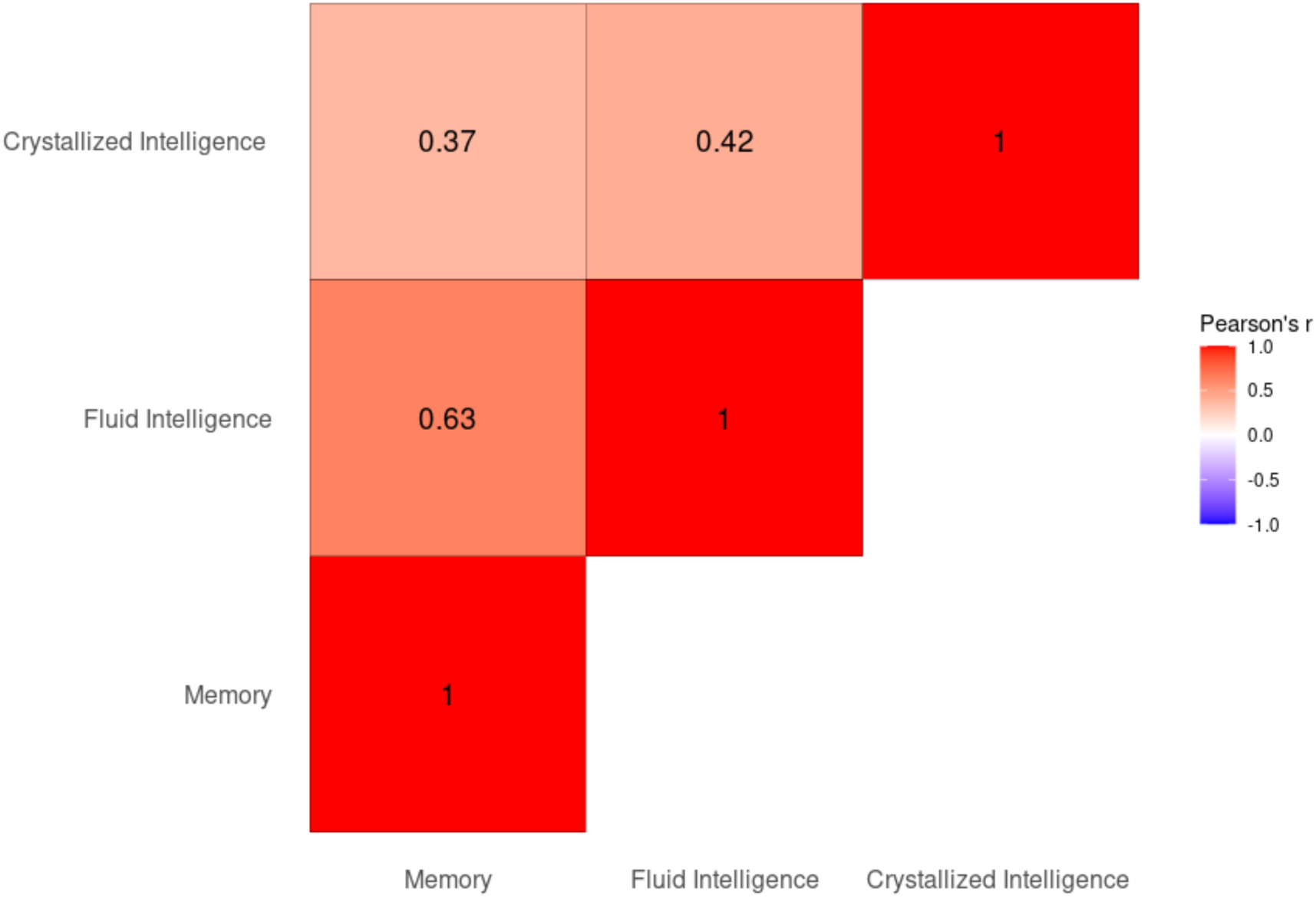
Pearson’s correlation of the three cognitive factor scores. Each factor appears to describe a different, but not entirely independent aspect of cognitive ability.

The first factor was weighted principally on the Rey Auditory Verbal Learning Task (RAVLT). The RAVLT is a widely used instrument for testing episodic memory. It consists of a list of 15 words (List A) that are read aloud to participants who then must immediately recall as many items as possible in 5 consecutive trials. Another list of 15 words (List B) is then presented, and participants are asked to immediately recall the new words. After the interfering list, participants are asked to recall words from List A ^43^. The first factor was therefore interpreted as a representation of the memory domain (memory).

The second factor was weighted principally on the NIH Toolbox ^44–46^ Dimensional Change Card Sort Test, Flanker Inhibitory Control and Attention Test, Pattern Comparison Test, and Trail Making Test (TMT) A & B. The Dimensional Change Card Sort Test is a test for complex attention and cognitive flexibility. Participants see a top card and two bottom cards. They are instructed to match one of the bottom cards to the top card according to different rules while inhibiting responses based on more salient and/or habitual rules. The Flanker Inhibitory Control and Attention Test is a test of inhibitory control and attention. It presents a target stimulus surrounded by four congruent or incongruent stimuli.

The participant must pay selective attention and respond to the target stimuli while suppressing responses to distractors. The Pattern Comparison Test is a test of processing speed. It presents two simple stimuli and participants are instructed to quickly respond whether the two stimuli are the same. Lastly, trail making test A & B is a test of complex attention and processing speed. TMT-A requires participants to connect randomly positioned circles following numeric order as quickly as possible, whereas TMT-B requires additional task switching ability as it requires participants to connect circles alternating between numeric order and alphabetic order. These four tests all tap into domains of processing speed, complex attention and cognitive flexibility. The second factor was therefore interpreted as a proxy for the fluid abilities (fluid intelligence).

The third factor was weighted principally on NIH Toolbox Picture Vocabulary Test and Oral Reading Recognition Test ^47^. The NIH Toolbox Picture Vocabulary Test is a receptive vocabulary test that does not involve either reading or writing. Participants are presented with a spoken word and then are asked to identify which picture out of four choices represents the word. The NIH Toolbox Oral Reading Recognition Test presents a word on a screen that participants then must identify out loud. These responses are then evaluated for pronunciation accuracy. Word reading and vocabulary tests are commonly used measures to gauge “premorbid function”, based on the premises that word reading abilities tap more into previously accumulated knowledge and is more resistant to age-related cognitive decline ^48^. We therefore interpreted the third factor as a proxy for crystalized abilities (crystalized intelligence). As illustrated in Figure 2, both the memory and fluid intelligence factors exhibit a strong correlation with age (memory: F=284.79, p < .01; fluid intelligence: F=613.08, p < .01), as anticipated, whereas crystalized intelligence factor was not significantly age-dependent (F=2.25, p = .1). A significant main effect of sex was found in memory performance (women performing better than men, F=82.15, p < .01) but not in other cognitive domains (fluid intelligence: F=2.82, p = .1; crystalized intelligence, F=.90, p = .34). Significant age-sex interaction was also found in memory (F=4.48, p = .03) and fluid intelligence performance (F=4.71, p = .03).

**Figure 2.**
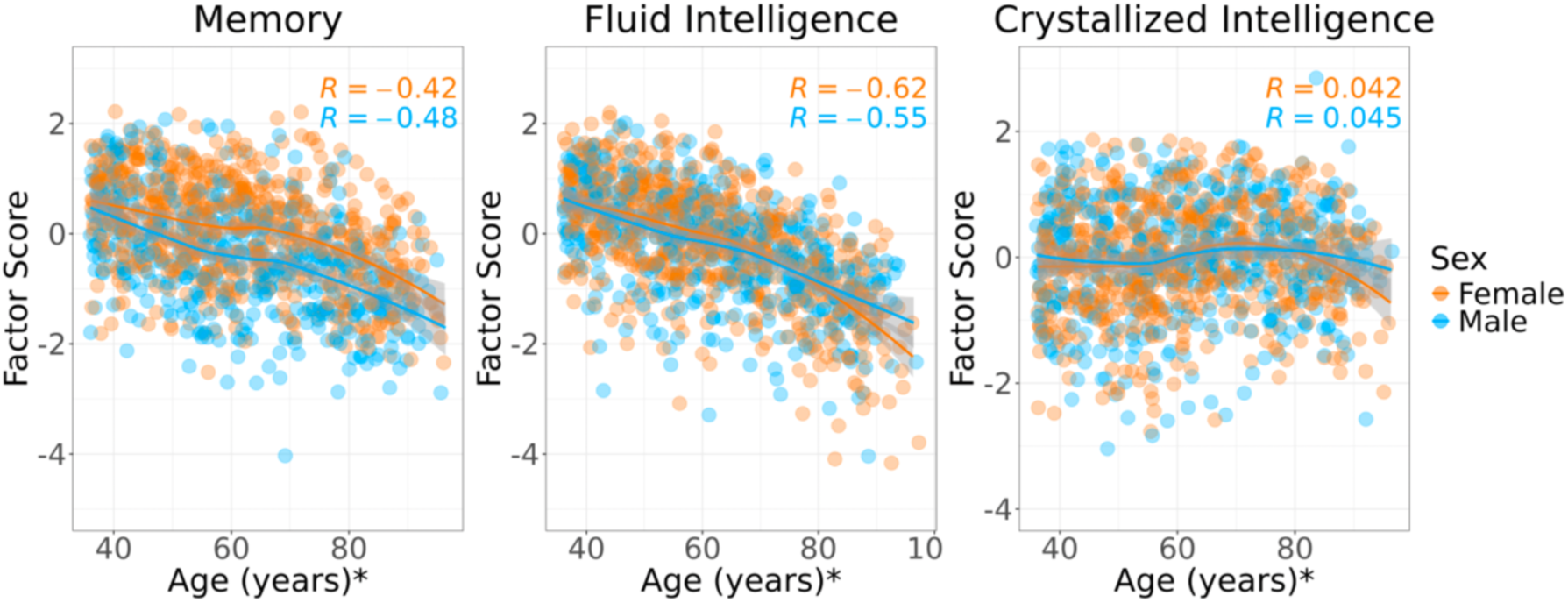
Associations with age for each factor score by sex. *The true age for the participants older than 90 years has been jittered on the figure for protection of personally identifiable information.

Performance shows substantial inter-individual variability across all three cognitive domains (Figure 2). Variance is largely age-stable in memory (χ²₁ = 0.05, p = .84) and fluid intelligence (χ²₁ = 0.23, p = .63) but decreases with age in crystallized intelligence (χ²₁ = 3.86, p = .050).

### 2.3 Performance groupings

Given the significant age and sex effects on cognitive performance, each individual’s residual value from a factor score-vs-age regression was used to classify their cognitive performance grouping (Figure 3). The regression lines were calculated within-sex for each decade of life to ensure that residuals accurately reflected performance relative to individuals with similar demographics and to account for non-linear effect of age on cognition. “Top performers” and “bottom performers” were defined as individuals in the top 25% of *positive* residuals and the bottom 25% of *negative* residuals, respectively, within their sex-stratified age bin. Secondary analyses grouped the individuals using the top and bottom 50% and 10% of residual values above and below the regression line, respectively, to examine how extremity of differences contributed to overall effects. Notably, top-performers had higher level of education compared to middle- and bottom-performers (Figure S1). Thus, this demographic difference was adjusted for in subsequent statistical analyses.

**Figure 3.**
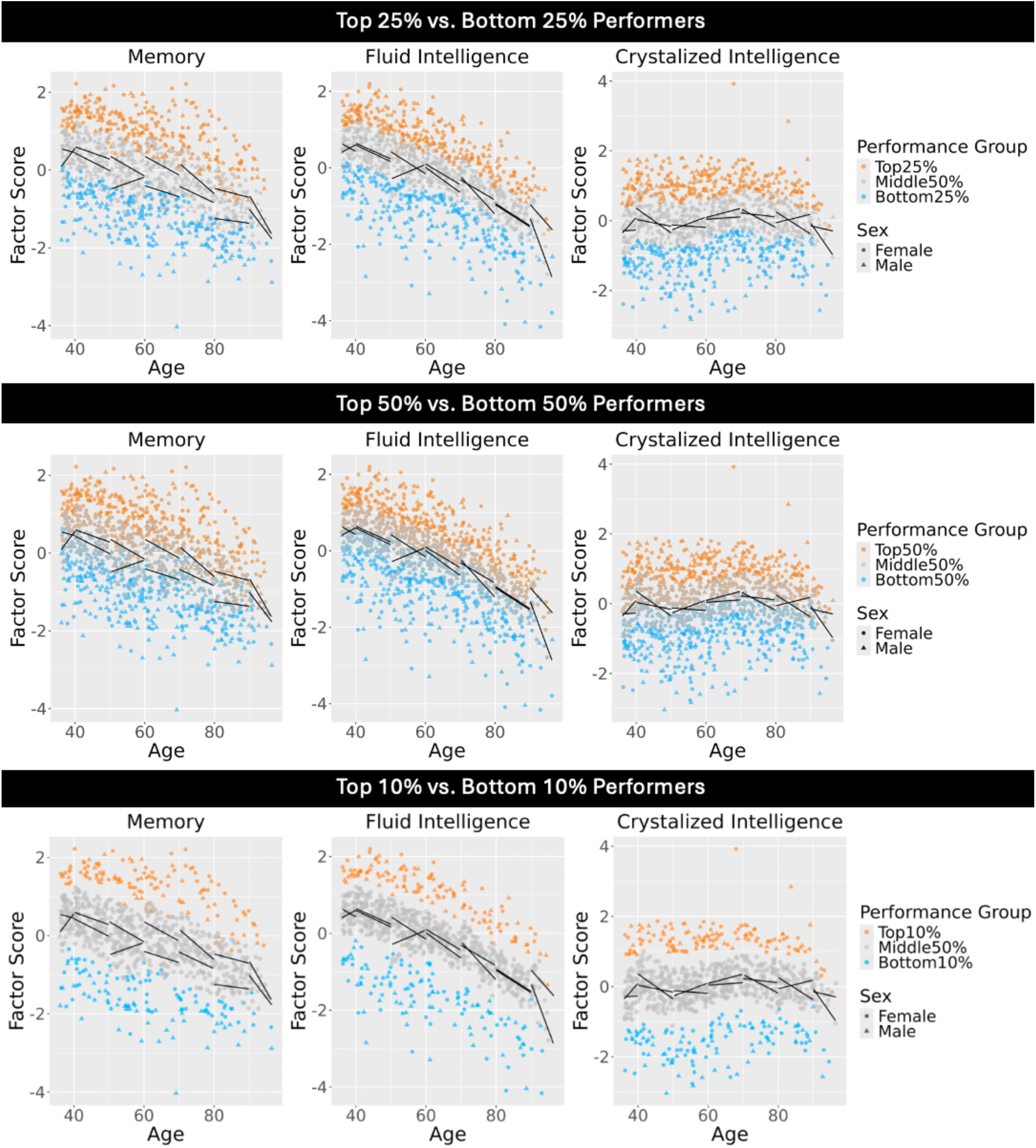
Generation of ‘top and ‘bottom’ cognitive performance groups. The upper row shows the distribution for the 25% groupings including the top quarter of positive residuals and the bottom quarter of negative residuals (more extreme differences in cognitive performance). The middle row shows the distribution for the 50% groupings including the top half of positive residuals and the bottom half of negative residuals (less extreme differences in cognitive performance). The bottom row shows the distribution for the 10% groupings including the top 10% of positive residuals and the bottom 10% of negative residuals (most extreme differences in cognitive performance). Note that when looking at the top percentile performers in the oldest participants, there is significant performance overlap with the middle 50% at much younger ages showing the superior performance in this sample. All groupings were performed within sex and age bin. Black lines show the regression lines for each sex and age bin. The ‘top’ group is shown in orange, the ‘bottom’ group is shown in blue, and the mid-range individuals are shown in gray. As can be seen in the plots, individuals in the top performance range in the latest decades of life (80+) perform similarly to the mean of the individuals in their 50s (this comparison to individuals 2-3 decades younger is typically used as a defining feature of a ‘superager’^11^), demonstrating the unique performance characteristics of this sample. This is particularly true for memory but less apparent for fluid intelligence which shows a steeper cross-sectional age decline in this sample where the 80+ group performs similarly to the low average of individuals in their 50s.

Initial analyses were performed in the total sample of participants (N=1146, 656 female). Additional analyses examined the continuous impact of age as well as effects within age bins including participants younger than 60 years (N= 560, 333 female), participants ranging from 60 to <80 years (N=390, 211 female), and participants 80 years and above (N=196, 112 female) to evaluate how the relationship between cognitive ability and cortical thickness may differ across cross-sectional age groups (Table 2). Because all groups were generated within sex and age bin independently, all performance groups have a proportionate sex and age distribution to the sample as a whole. More information about the cognitive performance groupings, including racial and ethnic demographics, APOE genotype, and years of education is available in Table S2.

**Table 2.** Cognitive scores in top and bottom performers by sex and age bin.

| **F1 (Memory)** |  |  |  |  |  |  |
| --- | --- | --- | --- | --- | --- | --- |
| **Age bin** | **Top 25%**<br>Average score (SD)<br>N (mean age)<br>N Females (mean age)<br>N Males (mean age) | **Bottom 25%**<br>Average score (SD)<br>N (mean age)<br>N Females (mean age)<br>N Males (mean age) | **Top 50%**<br>Average score (SD)<br>N (mean age)<br>N Females (mean age)<br>N Males (mean age) | **Bottom 50%**<br>Average score (SD)<br>N (mean age)<br>N Females (mean age)<br>N Males (mean age) | **Top 10%**<br>Average score (SD)<br>N (mean age)<br>N Females (mean age)<br>N Males (mean age) | **Bottom 10%**<br>Average score (SD)<br>N (mean age)<br>N Females (mean age)<br>N Males (mean age) |
| 36 – 90+ | .93 (.63)<br>283 (61.65)<br>163 (61.09)<br>120 (62.40) | -1.28 (.64)<br>280 (61.70)<br>160 (61.20)<br>120 (62.36) | 0.53 (.73)<br>580 (61.78)<br>332 (61.30)<br>248 (62.41) | -.85 (.73)<br>566 (61.55)<br>324 (61.05)<br>242 (62.22) | 1.23 (.59)<br>124 (62.06)<br>71 (61.77)<br>53 (62.47) | -1.63 (.61)<br>121 (61.60)<br>68 (61.03)<br>53 (62.33) |
| 36 – <60 | 1.28 (.39)<br>138 (48.15)<br>83 (48.28)<br>55 (47.95) | -.96 (.55)<br>137 (48.10)<br>82 (48.25)<br>55 (47.87) | .91 (.52)<br>281 (47.84)<br>167 (48.07)<br>114 (47.51) | -.52 (.64)<br>279 (47.82)<br>166 (47.92)<br>113 (47.67) | 1.55 (.30)<br>60 (48.29)<br>35 (48.03)<br>25 (48.66) | -1.34 (.51)<br>59 (47.70)<br>34 (47.48)<br>25 (47.99) |
| 60 – <80 | 0.86 (.52)<br>98 (69.00)<br>53 (68.41)<br>45 (69.70) | -1.43 (.59)<br>96 (69.26)<br>51 (68.85)<br>45 (69.72) | 0.44 (.62)<br>199 (69.26)<br>108 (68.77)<br>91 (69.83) | -.99 (.67)<br>191 (69.31)<br>103 (68.82)<br>88 (69.89) | 1.27 (.44)<br>42 (68.96)<br>23 (68.51)<br>19 (69.50) | -1.79 (.62)<br>40 (68.66)<br>21 (67.46)<br>19 (69.99) |
| 80 – 90+ | 0.04 (.50)<br>47 (85.93)<br>27 (86.06)<br>20 (85.75) | -1.91 (.40)<br>47 (85.89)<br>27 (86.06)<br>20 (85.66) | -0.36 (.61)<br>100 (86.05)<br>57 (85.94)<br>43 (86.20) | -1.53 (.53)<br>96 (85.99)<br>55 (86.10)<br>41 (85.85) | .30 (.48)<br>22 (86.46)<br>13 (86.82)<br>9 (85.94) | -2.12 (.36)<br>22 (86.04)<br>13 (86.07)<br>9 (85.99) |
| **F2 (Executive Functioning)** |  |  |  |  |  |  |
|  | **Top 25%** | **Bottom 25%** | **Top 50%** | **Bottom 50%** | **Top 10%** | **Bottom 10%** |

| Age bin | Average score | Average score | Average score | Average score | Average score (SD) | Average score (SD) |
| --- | --- | --- | --- | --- | --- | --- |
|  | N (mean age)<br>N Females (mean age)<br>N Males (mean age) | N (mean age)<br>N Females (mean age)<br>N Males (mean age) | N (mean age)<br>N Females (mean age)<br>N Males (mean age) | N (mean age)<br>N Females (mean age)<br>N Males (mean age) | N (mean age)<br>N Females (mean age)<br>N Males (mean age) | N (mean age)<br>N Females (mean age)<br>N Males (mean age) |
| 36 – 90+ | 0.74 (.71)<br>283 (61.73)<br>163 (61.30)<br>120 (62.32) | -1.32 (.84)<br>280 (61.62)<br>161 (61.01)<br>119 (62.45) | 0.41 (.76)<br>579 (61.73)<br>332 (61.39)<br>247 (62.20) | -.85 (.88)<br>567 (61.59)<br>324 (60.96)<br>243 (62.43) | 1.03 (.71)<br>124 (61.80)<br>71 (61.58)<br>53 (62.10) | -1.77 (.83)<br>122 (61.62)<br>69 (61.25)<br>53 (62.10) |
| 36 – <60 | 1.25 (.38)<br>137 (47.89)<br>82 (48.18)<br>55 (47.45) | -.83 (.61)<br>138 (47.95)<br>83 (48.03)<br>55 (47.82) | 0.93 (.46)<br>281 (47.72)<br>166 (47.95)<br>115 (47.40) | -.37 (.66)<br>279 (47.94)<br>167 (48.04)<br>112 (47.79) | 1.54 (.29)<br>60 (47.55)<br>35 (47.64)<br>25 (47.42) | -1.28 (.60)<br>60 (47.71)<br>35 (47.93)<br>25 (47.40) |
| 60 – <80 | 0.54 (.47)<br>99 (69.26)<br>54 (68.88)<br>45 (69.71) | -1.47 (.59)<br>95 (69.47)<br>51 (68.98)<br>44 (70.03) | 0.22 (.51)<br>198 (69.28)<br>109 (68.96)<br>89 (69.68) | -1.02 (.65)<br>192 (69.29)<br>102 (68.62)<br>90 (70.03) | .86 (.46)<br>42 (69.17)<br>23 (68.91)<br>19 (69.49) | -1.88 (.59)<br>40 (68.67)<br>21 (67.88)<br>19 (69.55) |
| 80 – 90+ | -0.32 (.46)<br>47 (86.25)<br>27 (86.01)<br>20 (86.58) | -2.43 (.68)<br>47 (85.92)<br>27 (85.85)<br>20 (86.01) | -0.66 (.51)<br>100 (86.16)<br>57 (86.04)<br>43 (86.32) | -1.94 (.73)<br>96 (85.88)<br>55 (86.00)<br>41 (85.73) | -.06 (.48)<br>22 (86.62)<br>13 (86.15)<br>9 (87.29) | -2.86 (.64)<br>22 (86.73)<br>13 (86.39)<br>9 (87.23) |
| **F3 (Crystallized Intelligence)** |  |  |  |  |  |  |
| Age bin | Top 25% | Bottom 25% | Top 50% | Bottom 50% | Top 10% | Bottom 10% |
|  | Average score<br>N (mean age)<br>N Females | Average score<br>N (mean age)<br>N Females | Average score<br>N (mean age)<br>N Females | Average score<br>N (mean age)<br>N Females | Average score (SD)<br>N (mean age)<br>N Females | Average score (SD)<br>N (mean age)<br>N Females |

|  | (mean age)<br>N Males<br>(mean age) | (mean age)<br>N Males<br>(mean age) | (mean age)<br>N Males<br>(mean age) | (mean age)<br>N Males<br>(mean age) | (mean age)<br>N Males<br>(mean age) | (mean age)<br>N Males<br>(mean age) |
| --- | --- | --- | --- | --- | --- | --- |
| 36 –<br>90+ | 1.06 (.40)<br>283 (61.59)<br>164 (61.10)<br>119 (62.25) | -1.16 (.55)<br>280 (61.65)<br>160 (61.11)<br>120 (62.36) | .68 (.51)<br>577 (61.77)<br>332 (61.25)<br>245 (62.47) | -.70 (.62)<br>569 (61.56)<br>324 (61.11)<br>245 (62.16) | 1.34 (.42)<br>124 (61.95)<br>71 (61.85)<br>53 (62.09) | -1.60 (.49)<br>123 (62.06)<br>70 (61.59)<br>53 (62.68) |
| 36 –<br><60 | 1.05 (.35)<br>138 (47.94)<br>83 (48.13)<br>55 (47.66) | -1.32 (.53)<br>138 (48.05)<br>83 (48.29)<br>55 (47.70) | .64 (.50)<br>281 (47.79)<br>167 (47.89)<br>114 (47.65) | -.84 (.64)<br>279 (47.87)<br>166 (48.10)<br>113 (47.53) | 1.33 (.28)<br>60 (48.23)<br>35 (48.46)<br>25 (47.90) | -1.79 (.46)<br>60 (47.96)<br>35 (48.08)<br>25 (47.79) |
| 60 –<br><80 | 1.16 (.41)<br>98 (69.07)<br>54 (68.61)<br>44 (69.64) | -.96 (.53)<br>95 (69.36)<br>50 (68.90)<br>45 (69.93) | .82 (.47)<br>196 (69.35)<br>108 (68.73)<br>88 (70.11) | -.52 (.59)<br>194 (69.22)<br>103 (68.87)<br>91 (69.62) | 1.46 (.44)<br>42 (68.75)<br>23 (68.40)<br>19 (69.17) | -1.42 (.47)<br>41 (69.39)<br>22 (68.44)<br>19 (70.48) |
| 80 –<br>90+ | 0.90 (.48)<br>47 (86.03)<br>27 (85.95)<br>20 (86.14) | -1.05 (.48)<br>47 (85.91)<br>27 (86.08)<br>20 (85.67) | 0.53 (.52)<br>100 (86.19)<br>57 (86.22)<br>43 (86.15) | -.66 (.53)<br>96 (85.85)<br>55 (85.81)<br>41 (85.91) | 1.11 (.59)<br>22 (86.41)<br>13 (86.31)<br>9 (86.56) | -1.40 (.46)<br>22 (86.88)<br>13 (86.40)<br>9 (87.56) |

Given the unpredictable nature of factors that may impact day to day cognitive performance, cognitive classification based on a single cross-sectional timepoint provides only partial information about an individual’s true cognitive capacity and limits the ability to classify as ‘high’ or ‘low’ performing. We therefore performed a set of secondary analyses to test the robustness of the baseline (V1) cognitive grouping implemented here. Specifically, we repeated the classification procedure in a longitudinal subsample (N = 657; 57.3%) with follow-up data collected an average of 2.67 years later (SD = 1.37). The same factor structure from the baseline sample (ages 36–80) was applied to the follow-up visit (V2) to generate factor scores. At V2, cognitive scores were residualized by both age and time between visits, stratified by sex and decade of life (using the same approach as the baseline timepoint). Individuals were classified as top or bottom performers based on percentile thresholds (top/bottom 25% or 50%) within their age–sex bin. Those who maintained the same 25 percentile grouping across V1 and V2 were labeled as *stable top 25%* or *stable bottom 25%* performers. This resulted in a substantially smaller subsample (memory: N=180, fluid intelligence: N=213, crystalized intelligence: N=226) but offered a conservative test of the robustness of our cortical brain structural results because analyses were limited to those who exhibited reliable cognitive grouping across sessions. Note, we did not aim here to assess longitudinal effects, but only to use the longitudinal stability as an indicator of grouping confidence. Additionally, we used a probabilistic approach to identify residual score ranges at V1 where individuals were most likely to switch group status at the 50% cutoff at V2. We estimated the likelihood of crossing the regression line from V1 to V2 and defined cutoff bands (e.g., -0.50 to 0.71 for memory residuals) where this probability was highest. Applying these bands back to the full baseline sample yielded high-confidence subsets of individuals whose 50% group classifications were most likely to hold longitudinally for each cognitive factor, labeled as *stable top 50%* or *stable bottom 50%* performers. Although these analyses do not provide the cognitive performance distinction that the more extreme groupings provide (i.e., the 25% and 10% groupings), they do ensure that the group label is likely to be stable across the full baseline sample. Generally, these secondary robustness analyses (conservative longitudinal and stable grouping samples) supported the overall conclusions derived from the full sample groupings reported here and therefore provide additional confidence in these results.

### 2.4 MRI data acquisition

Multimodal MRI data were acquired at each of the four HCP-A/AABC acquisition sites using a 3.0 Tesla Siemens Prisma scanner and 32-channel receive coils^42^. For this work, we used cortical thickness computed by FreeSurfer as a part of the HCP Pipelines using T1-weighted and T2-weighted images. The T1-weighted images were acquired using multi-echo magnetization-prepared rapid gradient-echo (MPRAGE) with prospective navigator motion correction (TR = 2500 ms; TI = 1000 ms; TE = 1.8/3.6/5.4/7.2 ms; 0.8 mm^3^ isotropic resolution; number of echoes = 4). T2-weighted SPACE images were also acquired with prospective navigator motion correction (TR = 3200 ms; TE = 564 ms, 0.8 mm^3^ isotropic resolution). More details on the imaging sequences have been described previously ^42^.

### 2.5 Areal feature-based parcellations

The structural data were processed by the AABC Informatics, Data Analysis, and Statistics Core (IDASC) using the HCP processing pipelines ^49,50^. Only the first two echoes of the T1-weighted MPRAGE sequence were used in analyses. Cortical surfaces were generated using both T1w and T2w images to improve the quality of cortical surface reconstructions (REF 48), and cortical thickness was computed as the distance between corresponding vertices on the white and pial surfaces. Areal feature-based (‘MSMAll’) registration^51^ was used to detect and align a complete cortical parcellation of 180 multi-modally defined cortical areas of interest as described in^50^. Detailed information about each of the mentioned cortical areas can be found in^50^. Cortical thickness estimates for each IDP were residualized on age within sex- and decade-based age bins before being included in regression analyses and cognitive performance group comparisons.

### 2.6 Statistical testing

The statistical modeling used in this study involved two different approaches: simple linear regression models to examine the association between each factor score and cortical thickness within each cortical area of the HCPMMPv1.0 parcellation, and one-way ANCOVA models to test for group differences in mean cortical thickness within each cortical area between top and bottom performers (e.g., top/bottom 25% performers). Additional analyses were performed to examine these relationships by sex and to compare top-/bottom-performers with general population (e.g., mid 50% performers). All statistical tests of imaging data were performed in each hemisphere independently and were conducted using R^52^. Linear models were generated using the Ordinary Least Squares regression method, and one-way ANCOVA tests were conducted using the aov() function. Age, sex, and years of education were included as covariates in all models, except for those conducted within each sex group. Robustness tests were conducted to compare cognitive performance between the longitudinally defined stable 25% and stable 50% groups.

False Discovery Rate (FDR) correction was performed using Benjamini-Hochberg method to control for false positive results^53^ with the p.adjust() function in R^52^. The effect size of each ANCOVA model was measured by calculating Cohen’s d. This was performed by computing the difference between the mean cortical thickness of top and bottom performers and dividing that value by the pooled standard deviation of cortical thickness within each cortical area. Data visualization was performed in R^52^ and Connectome Workbench visualization software^54^.

## 3. Results

### 3.1 Associations between age and cortical thickness

Increasing age was associated with decreasing cortical thickness in the entire sample using a surface-based simple linear model controlling for sex. The results suggest global thinning with age, but particularly strong associations were observed in primary somatosensory regions, prefrontal cortex, and operculum regions (Figure 4). These results appear largely consistent with previous findings^24^.

**Figure 4.**
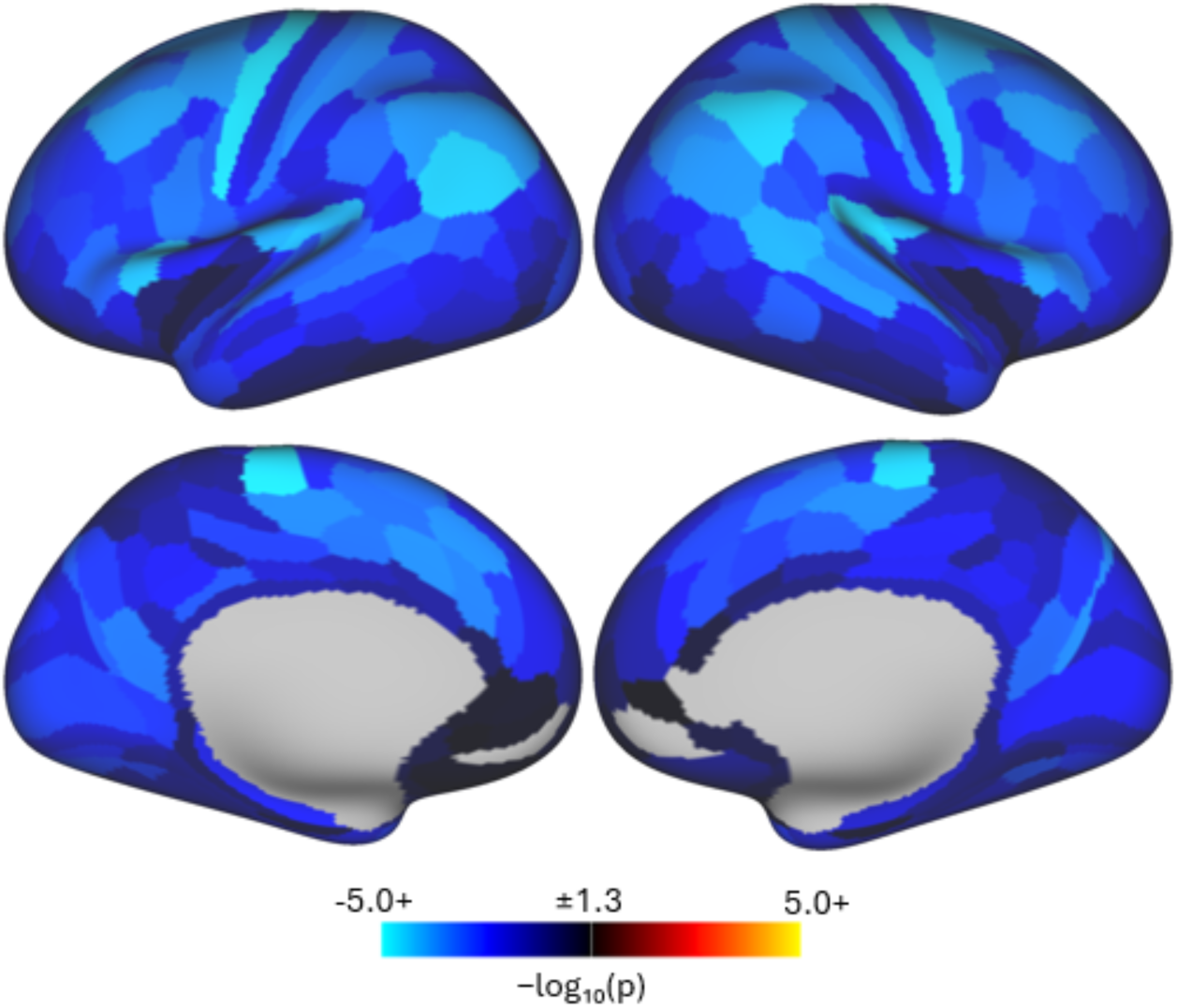
Spatial distribution of age effect on cortical thickness in our sample. Color scale shows signed −log₁₀(p) (value = −log₁₀(p) × sign[β]); positive values (warm colors) indicate cortical areas where older age is associated with greater cortical thickness, negative values (cool colors) indicate cortical areas where older age is associated with thinner cortex. Only cortical areas surviving FDR correction (p < 0.05; |−log₁₀(p)| > 1.3) are shown; non-significant areas are masked. Negative age–thickness associations remained significant after FDR correction in 356 of 360 cortical areas.

### 3.2 Associations between cognitive ability and cortical thickness

Higher factor score residuals (better cognitive performance after accounting for age and sex) were significantly associated with thicker cortex in the entire sample using cortical areal parcellations in simple linear models (Figure 5). Given the strong effects of education on cognitive performance (Figure S2), years of education was included as a covariate in all models. All analyses on cortical thickness and cognitive ability were visualized to show whole-cortex effect-size maps, displaying the standardized effect size for every cortical area. Areas that met statistical significance threshold are outlined in white. This presentation follows recent recommendations to report both unthresholded effect sizes and thresholded results in neuroimaging, moving beyond binary significance to provide complete, interpretable results^55^.

**Figure 5.**
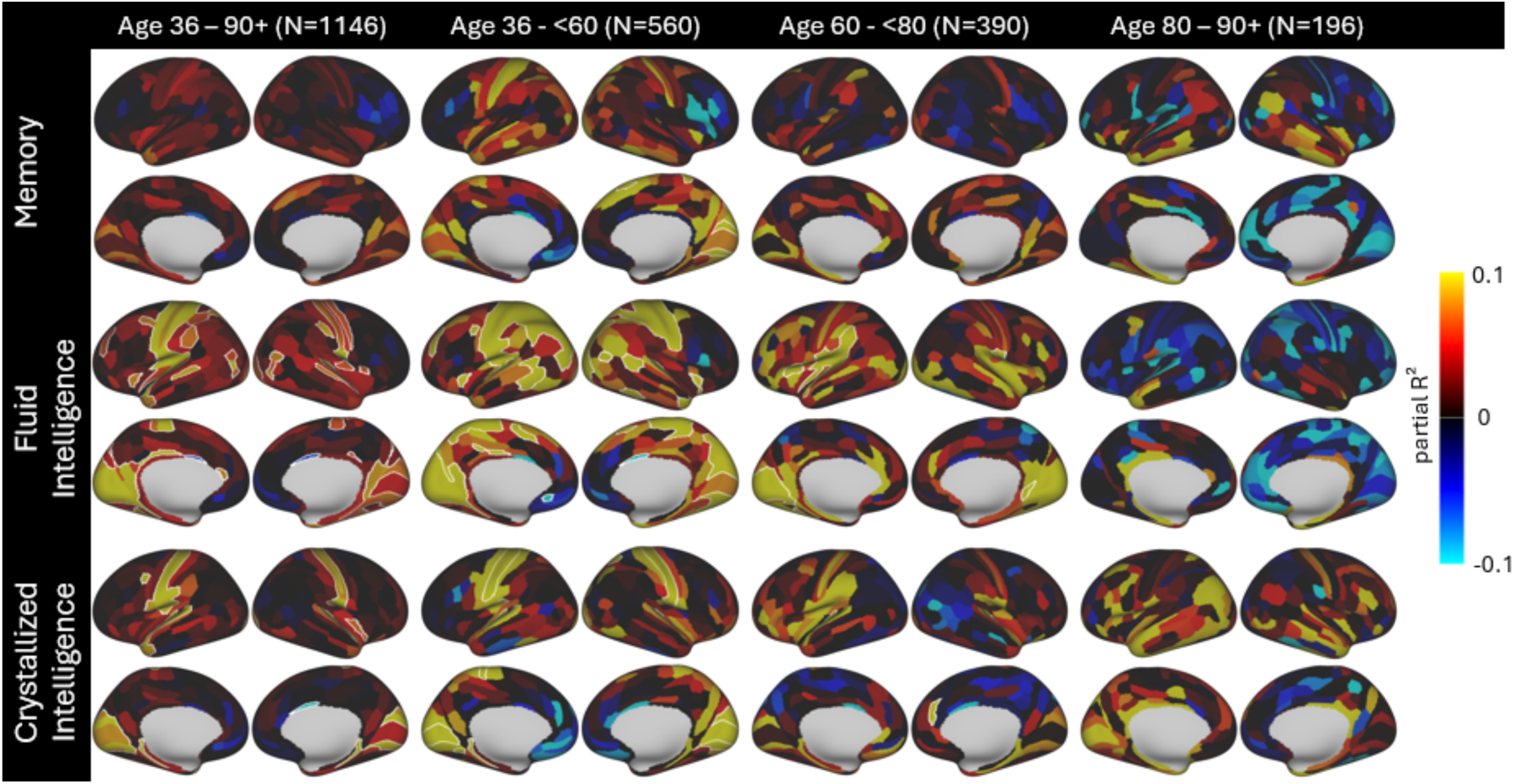
Spatial maps illustrate the associations between cognitive factor–score residuals and cortical-thickness residuals for each cortical area, after adjusting for age, sex, and education. Effect sizes are expressed as partial R² values and are signed according to the direction of association (warmer colors indicate cortical areas where higher cognitive scores are associated with greater cortical thickness; cooler colors indicate associations with thinner cortex). Cortical areas surviving FDR correction (p < 0.05) are outlined in white.

The fluid intelligence factor score exhibited stronger associations with cortical thickness than memory and crystallized intelligence scores as indicated by larger effect sizes overall (brighter colors) and more widespread cortical areas with statistical significance (outlined). The effects were more prominent in midlife adults (age 36 – <60) than older adults (age 60 – <80, and 80 – 90+ groups) and were dominant in primary and secondary sensory and motor areas across cerebral cortex. In the midlife adults group (age 36 – <60), the cortical areas with the highest significance were located in the somatomotor cortex (e.g., 1, 2, 4, 5m, 5L), visual cortex (e.g., V3A, V7, LIPv), auditory cortex (e.g., A1, MBelt), and intraparietal cortical areas (e.g., IP1, IP2, MIP). In the younger-old group (60 -<80), higher fluid ability remained strongly associated with thicker cortex in cortical areas of somatomotor cortex (e.g., OP1, OP2_3, OP4), auditory and visual processing (e.g., A1, Lbelt, V1, V2, VMV1), insular cortex (e.g., FOP3, FOP4), and one isolated cortical area in the medial temporal lobe (PHA3), demonstrating a partial independent replication of effects found in the younger sample. However, the overall number of cortical areas showing significant associations decreased compared to the younger group. These effects were not replicated in the older adults groups (80 – 90+), which also had notably smaller sample sizes impacting statistical power. However, as can be seen in Figure 5, the effect sizes in the older adult groups were also comparably smaller (darker color) and showed variable directionality (both positive and negative associations), suggesting that power alone does not explain these differences.

Similar to fluid intelligence scores, higher crystallized intelligence scores were associated with increasing cortical thickness largely in cortical areas related to sensory processing in the middle-aged (36 - <60) group, including primary auditory cortex (i.e., A1, MBelt, PBelt), visual cortex (i.e., V1, V2, V3, VMV1, LIPv), somatomotor cortex (e.g., area 1, 3a, 3b, OP1) and one cortical area in the medial temporal lobe (PreS). In the age 60 - 80 group, better crystallized intelligence performance was associated with thicker cortex in primary auditory cortex (A1, MBelt) and one cortical area in anterior cingulate cortex (area d32).

Comparably larger effect sizes were found in the operculum regions. No statistically significant effects were found in the age 80+ group, though subthreshold large effect sizes were found in regions of superior temporal lobe, inferior parietal lobule, medial occipital lobe, and cingulate cortex.

The memory factor score was found to have minimal association with cortical thickness throughout the age groups. Significant effects were found in a few isolated cortical areas in the middle-aged adult (36 - <60) group, including cortical areas of visual processing (V2, V6A, and V7), one cortical area in somatomotor cortex (5L) and a language area (SFL). No statistically significant results were found in the older adult groups (60 - 80 and 80 - 90+) after correction. Given the differences in spatial distribution of the effects observed in three age groups, we performed formal interaction tests and confirmed significant effects of the interaction between cognitive performance and age group on cortical thickness in certain cortical areas (Figure S3).

Secondary analyses examined the associations between cognitive performance and cortical thickness separately in men and women. The strongest effects were observed in midlife (36 - <60) females, where fluid intelligence was strongly associated with thicker cortex in the visual cortex, somotomotor cortex, auditory cortex, and intraparietal regions, largely mirroring the results from the combined-sex analysis (Figure 6). Memory was also found to relate with thicker cortex in those regions with similar effect sizes in midlife (36 - <60) women. The male group (a smaller sample), in contrast, showed bigger effect sizes in older groups (60 - 80 and 80 - 90+) when associating cortical thickness with fluid intelligence and crystalized intelligence. However, those effects did not reach threshold for statistical significance. The most prominent associations as measured by significance level in males were between crystallized intelligence and cortical thickness in the primary auditory cortex and somatomotor cortex, and these effects emerged only when using the full age-range sample (Figure 7).

**Figure 6.**
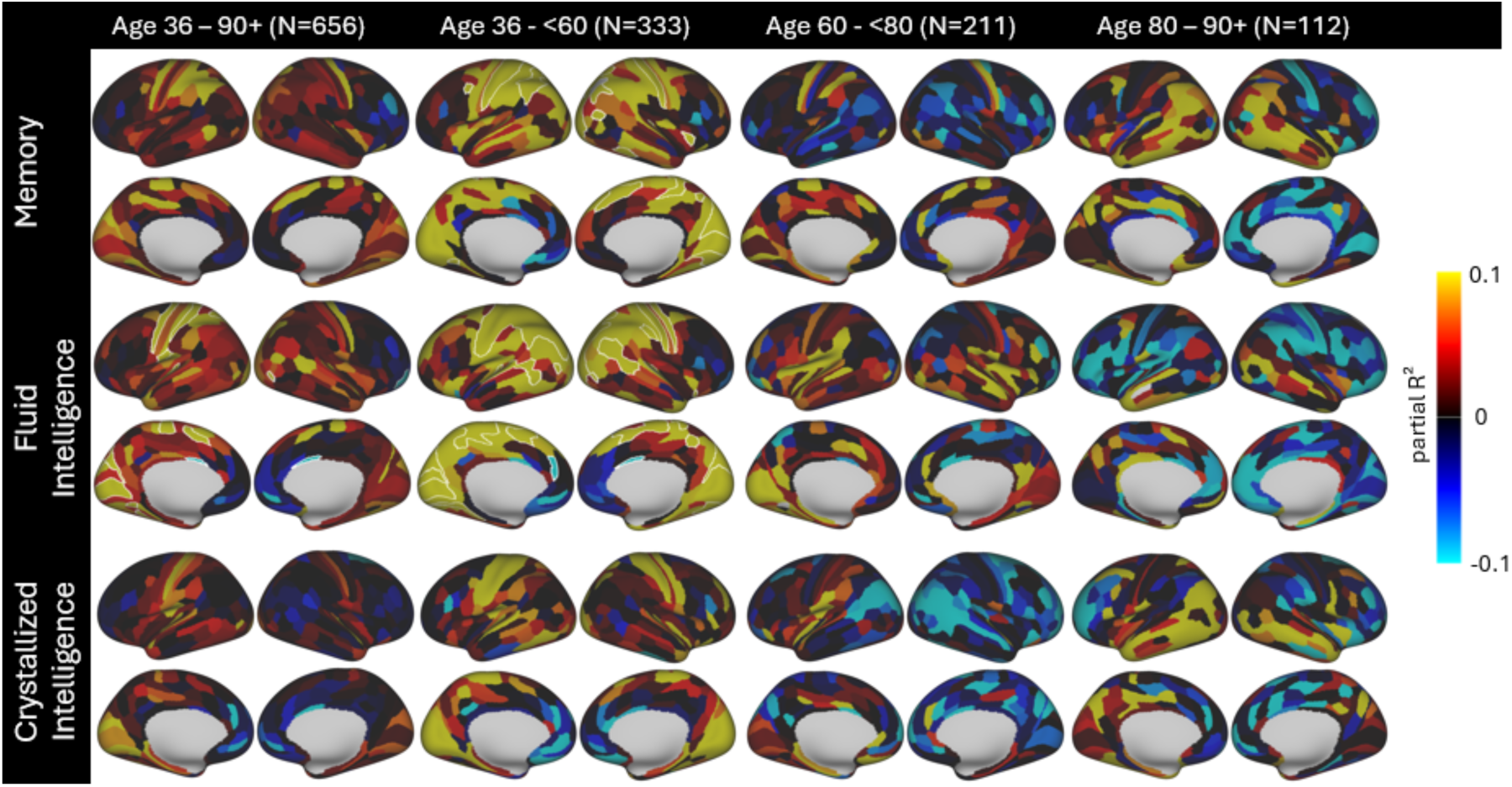
Spatial maps illustrate the associations between cognitive factor–score residuals and cortical-thickness residuals for each cortical area in *female* participants, after adjusting for age and education. Effect sizes are expressed as partial R² values and are signed according to the direction of association (warmer colors indicate cortical areas where higher cognitive scores are associated with greater cortical thickness; cooler colors indicate associations with thinner cortex). Cortical areas surviving FDR correction (p < 0.05) are outlined in white.

**Figure 7.**
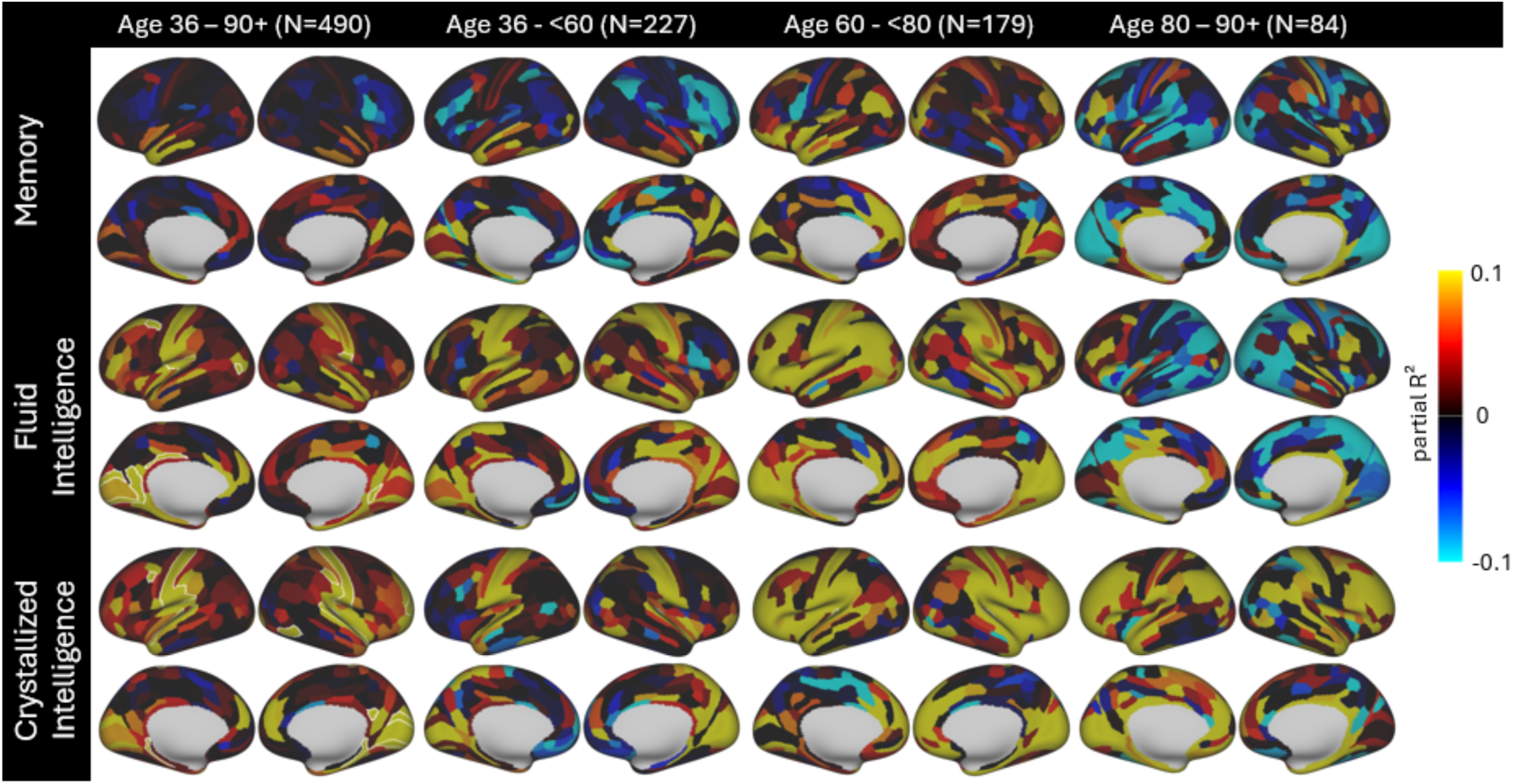
Spatial maps illustrate the associations between cognitive factor–score residuals and cortical-thickness residuals for each cortical area in *male* participants, after adjusting for age and education. Effect sizes are expressed as partial R² values and are signed according to the direction of association (warmer colors indicate cortical areas where higher cognitive scores are associated with greater cortical thickness; cooler colors indicate associations with thinner cortex). Cortical areas surviving FDR correction (p < 0.05) are outlined in white.

An additional comparison between males and females, using the full age-range sample and uncorrected p-values, revealed more similar spatial patterns across groups. Specifically, both sexes showed strong involvement of the somatomotor, visual, and auditory cortices in relation to performance on fluid intelligence tasks. However, male participants showed stronger associations between crystallized intelligence and cortical thickness, whereas female participants exhibited stronger effects in relation to memory (Figure 8). Overall, these effects suggest that an overlapping set of regions show covariation in thickness relative to cognitive performance in male and female participants.

**Figure 8.**
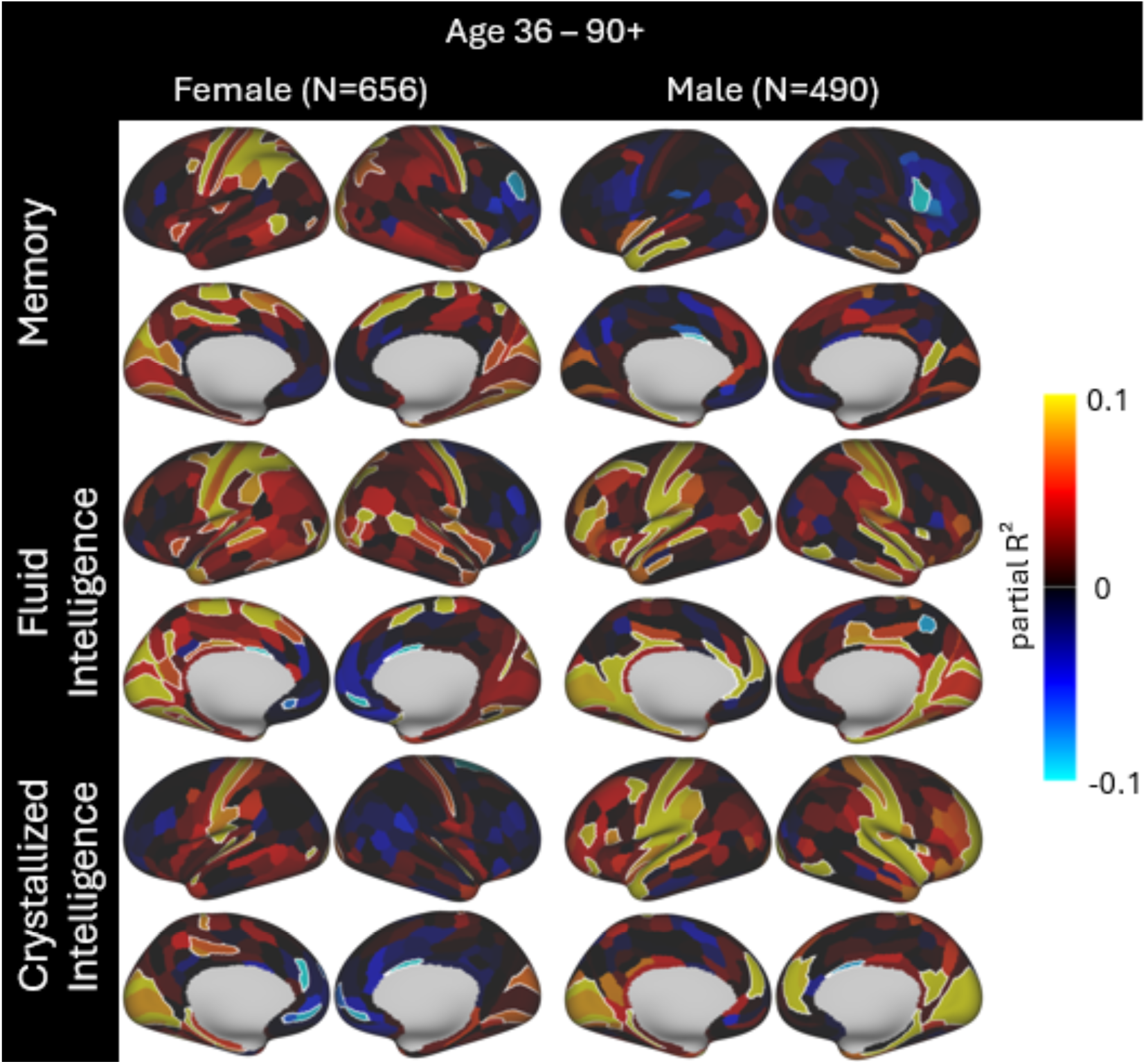
Comparison of the spatial distribution of associations between cognitive factor–score residuals and cortical-thickness residuals for each cortical area in the full age-range sample by sex, after adjusting for age and education. Effect sizes are expressed as partial R² values and are signed according to the direction of association (warmer colors indicate cortical areas where higher cognitive scores are associated with greater cortical thickness; cooler colors indicate associations with thinner cortex). Significant cortical areas (uncorrected p < 0.05) are outlined in white.

Specifically, the memory factor score residual was associated with one cortical area in a higher-order visual processing area (POS1) in both men and women. The fluid intelligence factor score residual was associated with cortical areas in somatomotor cortex (e.g., 1, 3a, 3b), visual-related cortices (e.g., MT, V1, VMV1, POS1), auditory cortex (e.g., A1, MBelt, TA2), posterior cingulate cortex (23d, d23b), and a parahippocampal cortical area (PHA1) in both sex groups. The crystallized intelligence factor score residual was associated with cortical areas in somatomotor cortex (e.g., 1, 3b, OP1), visual cortices (e.g., V1, V2, VMV1), auditory cortex (e.g., A1, A4, MBelt), and a MTL cortical area (PreS) in both men and women. It may be appropriate to consider the uncorrected effects given that the analysis in part is an independent replication of effects in two distinct cohorts (male and female). Given this, these sex-stratified analyses provide additional confidence of the importance of the set of identified replicated regions in supporting cognitive performance. However, effects differ in effect size, the full regional distribution, and the cognitive domain and age-sample showing the strongest associations. Male participants also exhibited reduced lateralization, with more similar left–right spatial distributions; whereas female participants appeared to show a greater level of left-hemisphere lateralization. Any differences may be due to true sex differences but could also be related to variations in sample size and statistical factors as well as other factors that are not matched between the groups. We primarily aimed to examine major influences of sex on the main results here and more formal analyses of sex effects as well as factors that contribute to sex differences will be performed in future work.

### 3.3 Group differences in cortical thickness between top and bottom 25% performers

Compared to the full sample association analyses demonstrated in Figure 5, similar regions were implicated in the ANCOVA analyses evaluating group differences in cortical thickness between top 25% and bottom 25% cognitive performers controlling for age, sex and education (Figure 9). Strongest effects were observed between fluid intelligence performance and cortical thickness in the midlife (36 - <60) adult group, with strongest involvement of visual cortices (e.g., LIPv, V3A, V3) and somatomotor cortex (e.g., 3a, 3b, 4, 5m). In the same age group, crystalized intelligence score revealed a similar pattern involving visual cortices (e.g., VMV1, V2, V3), somatomotor cortex (e.g., 3a, 4, 5m) and auditory cortex (e.g., A1, MBelt, PBelt). The group with better memory performance in the midlife age range was found with mixed effects, including thicker cortex in a few isolated cortical areas (e.g., 5L, VMV1, PEF, A1, STSvp) and thinner cortex in a few cortical areas in the frontal cortex (44, IFSp, a10p, 33pr, 6r).

**Figure 9.**
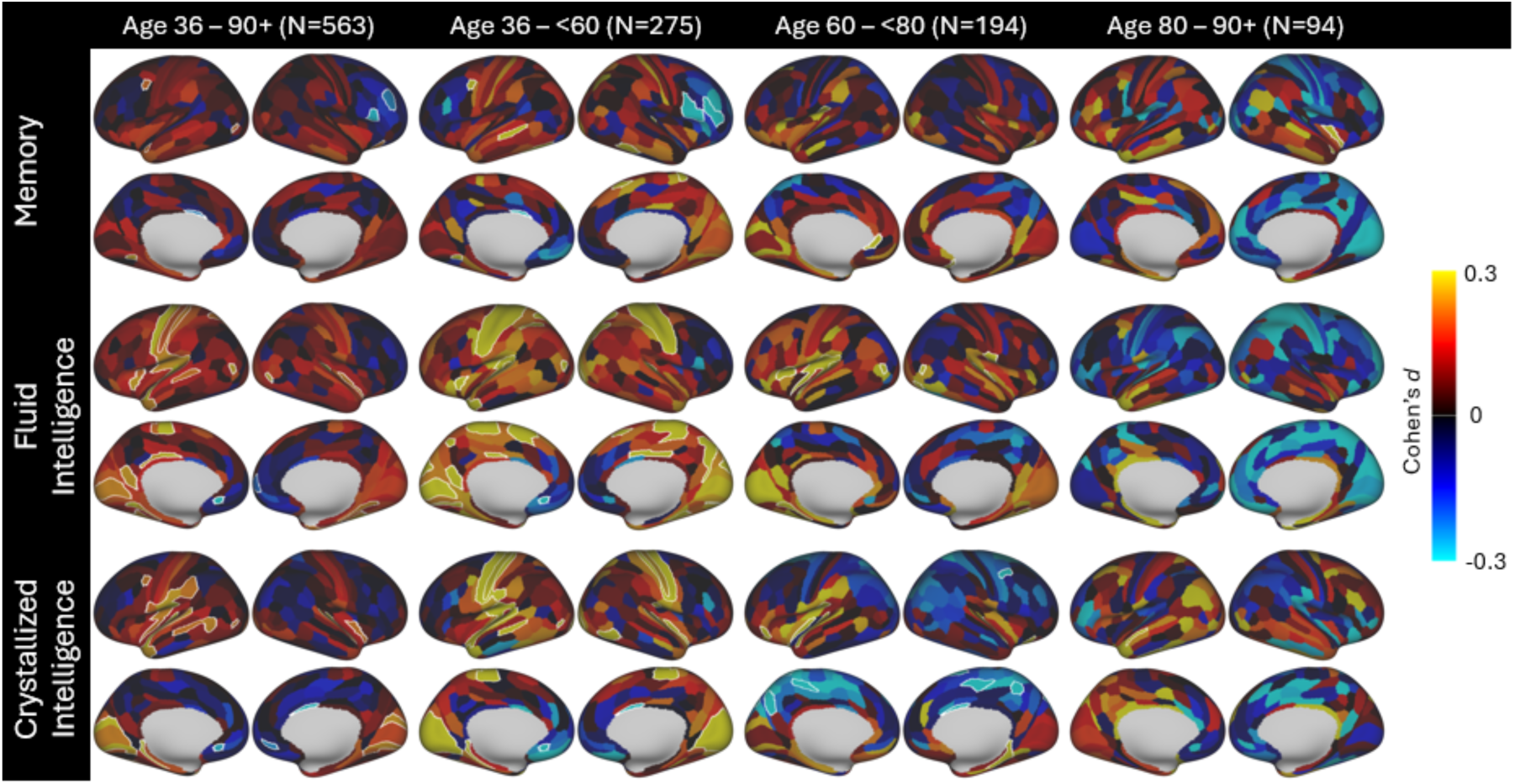
Spatial maps illustrate the group difference in cortical thickness residuals for each cortical area between top 25% and bottom 25% of cognitive performers, after adjusting for age, sex, and education. Effect sizes are expressed as Cohen’s *d* values and are signed according to the direction of group difference (warmer colors indicate cortical areas where cognitive high performers show greater cortical thickness; cooler colors indicate high performers with thinner cortex). Cortical areas surviving FDR correction (p < 0.05) are outlined in white.

In the young-old group (60 – <80 years), group differences between top and bottom performers were less prominent overall but showed significant involvement of visual, auditory, somatomotor cortices and regions of frontal operculum on fluid intelligence performance. Cortical thickness comparison on crystalized intelligence performance grouping in this age range became mixed, with thicker cortex found in isolated cortical areas in auditory cortex and medial temporal lobe (PHA1, PreS) while thinner cortex found in isolated cortical areas in visual cortices, somatomotor cortex and parietal lobe. Better memory performance was found with thicker cortex in a small number of isolated cortical areas (a24, 47s, MBelt, OP1, pOFC).

In the older group (80 – 90+), with smaller sample sizes and lower statistical power, only a few isolated cortical areas remained statistically significant after correction. Specifically, in the older adult group (80 – 90+), fluid abilities were associated with thicker cortex in perirhinal entorhinal cortex (PeEc), crystalized abilities were associated with thicker cortex in a language processing cortical area (STSda) and temporal fusiform cortex (TF), and the memory factor score was associated with thicker cortex in another language processing cortical area (STGa) and a cortical area in insular cortex (Pol2). Similar to the association analyses findings, the older adult groups also exhibited a mix of positive and negative effects compared to younger groups.

Contrary to expectations that elevated cortical thickness would be a unique feature of high performers in this generally healthy sample (i.e. as a mechanism of cognitive resilience), closer examination of regional effects demonstrated that group differences in cortical thickness were due to the low performers (although still cognitively healthy) having reduced cortical thickness relative to the typically performing group (i.e., the middle 50% performing portion of the sample; Figure 10) in the abovementioned regions including somatomotor, auditory and visual cortices. The high performers showed generally smaller effect sizes with limited cortical areas showing significantly thicker cortex compared to the middle performing sample, and the findings are more directionally mixed (Figure 11). However, a few cortical areas were found with significant differences in favor of high performers in the oldest portion of the sample, including the left entorhinal cortex (L_EC) and cortical areas in posterior cingulate cortex (31pd) and superior temporal lobe (TA2) and may be of interest for future studies.

**Figure 10.**
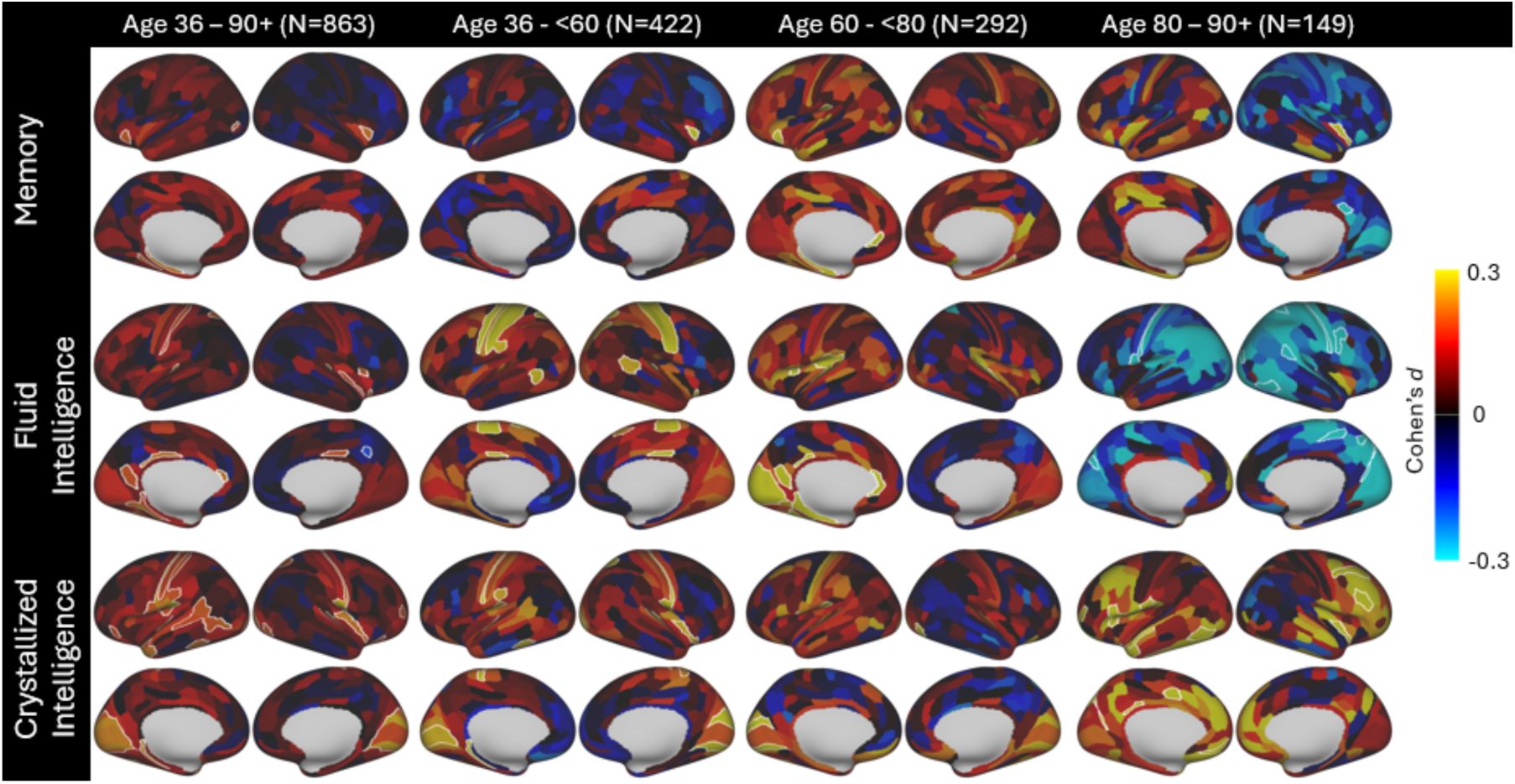
Spatial maps illustrate the group difference in cortical thickness residuals for each cortical area between bottom 25% and middle 50% of cognitive performers, after adjusting for age, sex, and education. Effect sizes are expressed as Cohen’s *d* values and are signed according to the direction of group difference (warmer colors indicate cortical areas where cognitive high performers show greater cortical thickness; cooler colors indicate high performers with thinner cortex). Cortical areas surviving FDR correction (p < 0.05) are outlined in white.

**Figure 11.**
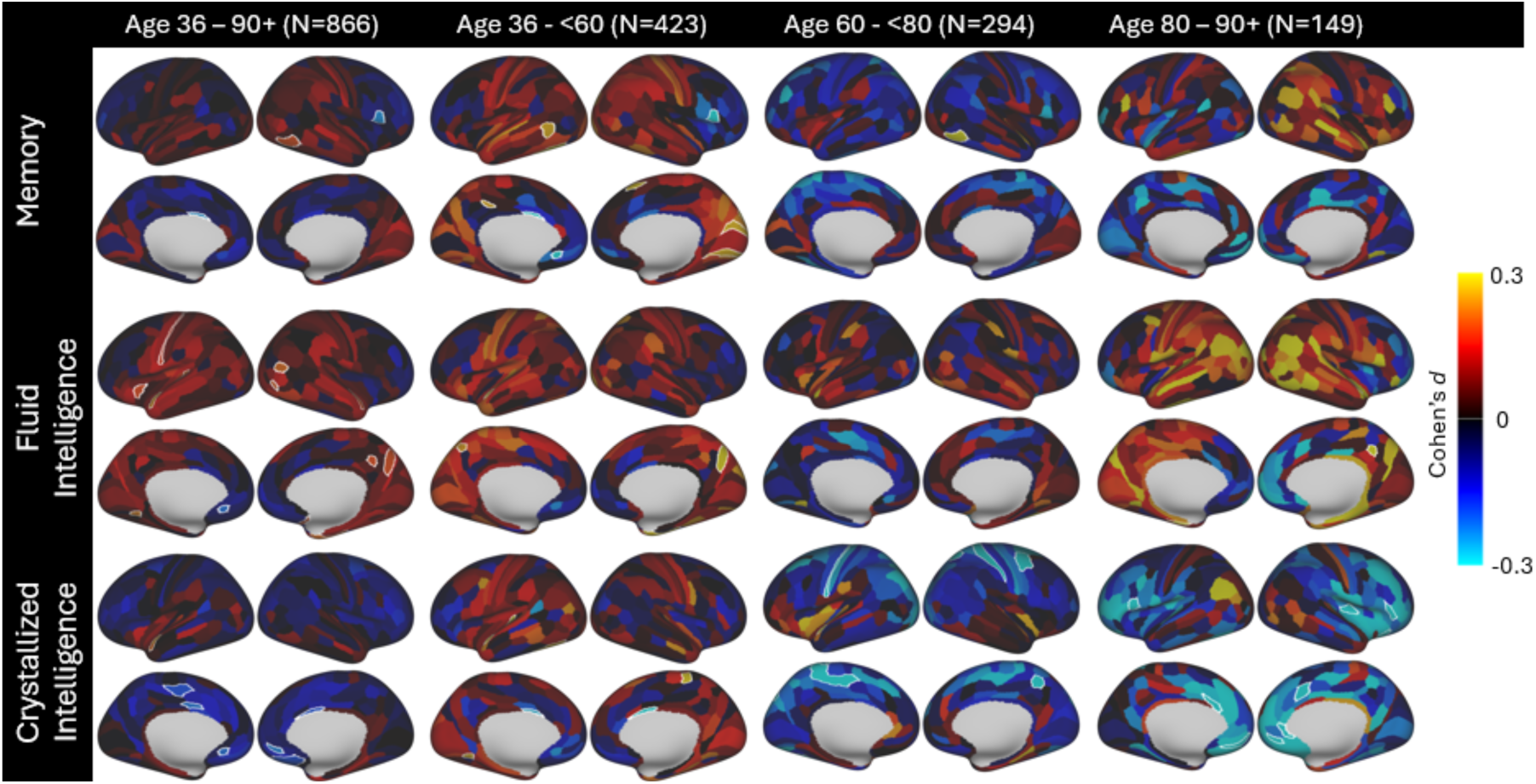
Spatial maps illustrate the group difference in cortical thickness residuals for each cortical area between top 25% and middle 50% of cognitive performers, after adjusting for age, sex, and education. Effect sizes are expressed as Cohen’s *d* values and are signed according to the direction of group difference (warmer colors indicate cortical areas where cognitive high performers show greater cortical thickness; cooler colors indicate high performers with thinner cortex). Cortical areas surviving FDR correction (p < 0.05) are outlined in white.

To better illustrate the age-related changes in cortical thickness across cognitive performance groups, and to help better understand some of the observed counterintuitive findings where high performers were found with thinner cortex, we generated single-area plots (Figure 12). We plotted three cortical areas in the somatomotor, auditory, and visual cortices, respectively, that showed the highest statistical significance with comparably larger effect sizes (Cohen’s *d* ≥ 0.5) and were reliably replicated across various subsample analyses. In these cortical areas, a convergence effect, and in some cases even a crossover, was observed, such that the differences in cortical thickness between cognitive performance groups diminished with increasing age. As age increases, the direction of the effect even reversed, with higher-performing individuals exhibiting thinner cortices. In some cortical areas, this cross over effect was strong enough to show a significant finding in our ANCOVA analyses where high-performing group show significantly thinner cortices in the 80+ age bin (Figure 13). Interaction tests support significant effects of the interaction between age and cognitive performance group on cortical thickness in some cortical areas (Figure S4). These patterns may suggest that the neural mechanisms supporting cognitive performance differ between younger and older adults, potentially reflecting age-dependent shifts in brain–behavior relationships. These results must be reviewed with caution given the cross-sectional nature of this study. They should also be viewed considering biological processes that can increase the apparent gray–white boundary distance produced by image-processing (e.g., tissue changes that alter contrast), potentially inflating thickness estimates.

**Figure 12.**
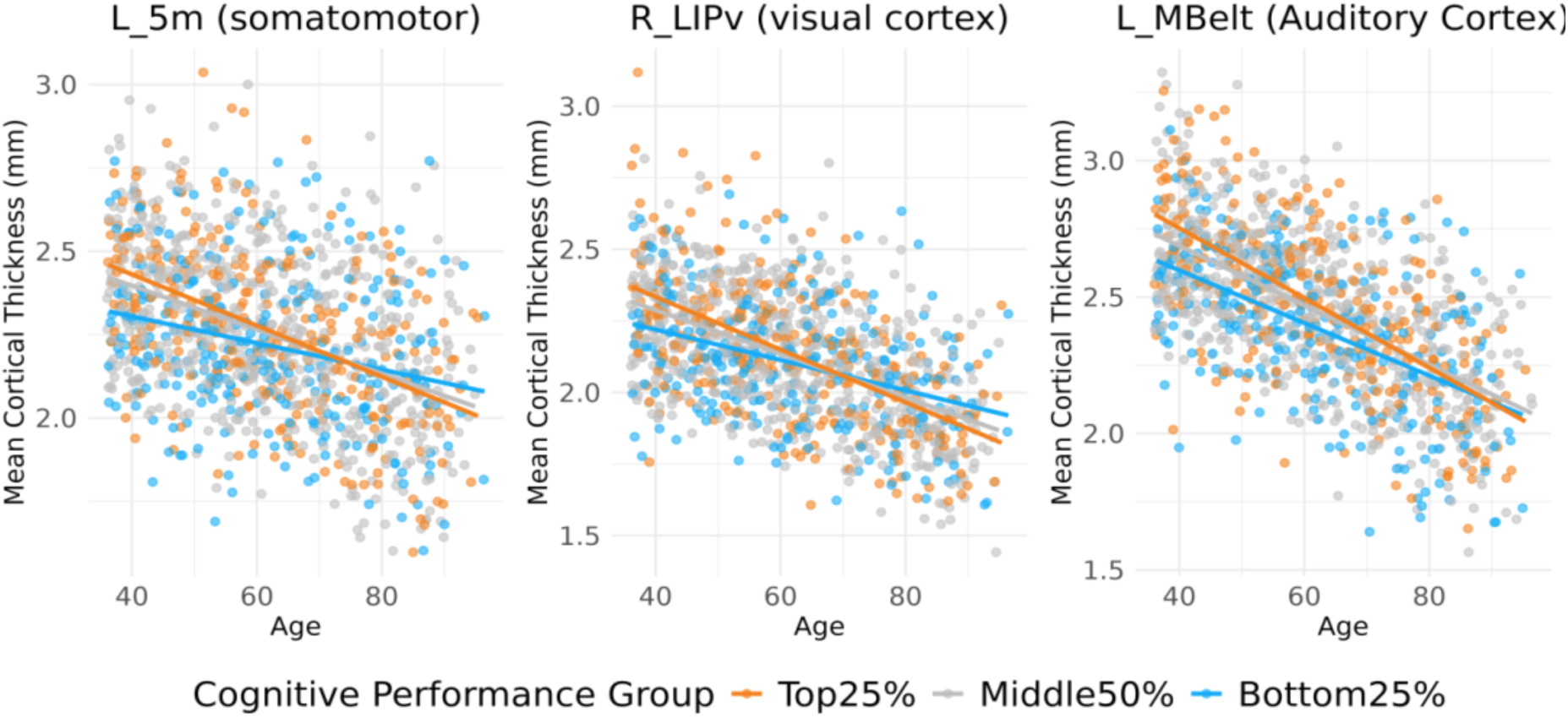
Scattered plots show group differences in cortical thickness decrease or cross over with age in three cortical areas in the somatomotor, auditory, and visual cortices, respectively. These areas were selected as they showed the highest statistical significance with comparably large effect sizes (Cohen’s *d* ≥ 0.5) and were reliably replicated across various subsample analyses. *The true age for the participants older than 90 years is jittered on the figure to protect personally identifiable information.

**Figure 13.**
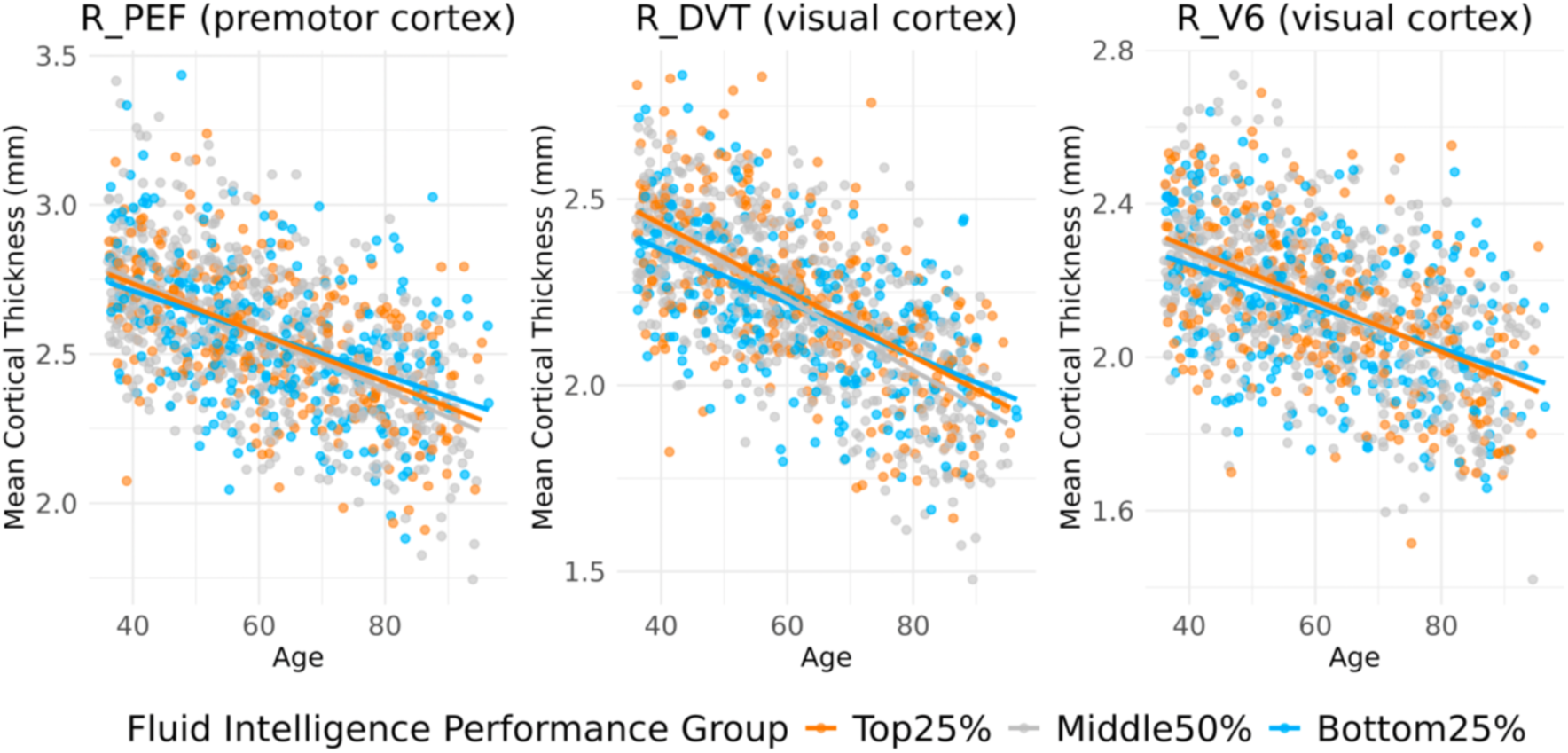
Scattered plots show group differences in cortical thickness cross over with age in cortical areas where we found thinner cortex in high- performers. *The true age for the participants older than 90 years is jittered on the figure to protect personally identifiable information.

Additional analyses were performed to examine robustness of findings by expanding the performance groupings to a broader middle range (i.e., the 50% splits) and, conversely, restricting them to the most extreme performers (i.e., the 10% splits). For contrast, both sets of grouping results are displayed on an identical effect-size scale (−0.5 to 0.5; see Figs. 14–15). Although the 10% grouping substantially reduced sample size, it yielded markedly larger effect sizes and a significantly greater number of suprathreshold cortical areas (Figure 14) relative to the median split (Figure 15). These findings suggest that the statistical significance in the ANCOVA comparisons is disproportionately driven by individuals at the extremes of cognitive performance. Although effect sizes varied, the spatial distribution of significant positive associations of the 50% and 10% grouping remained broadly consistent with the quartile-based results.

**Figure 14.**
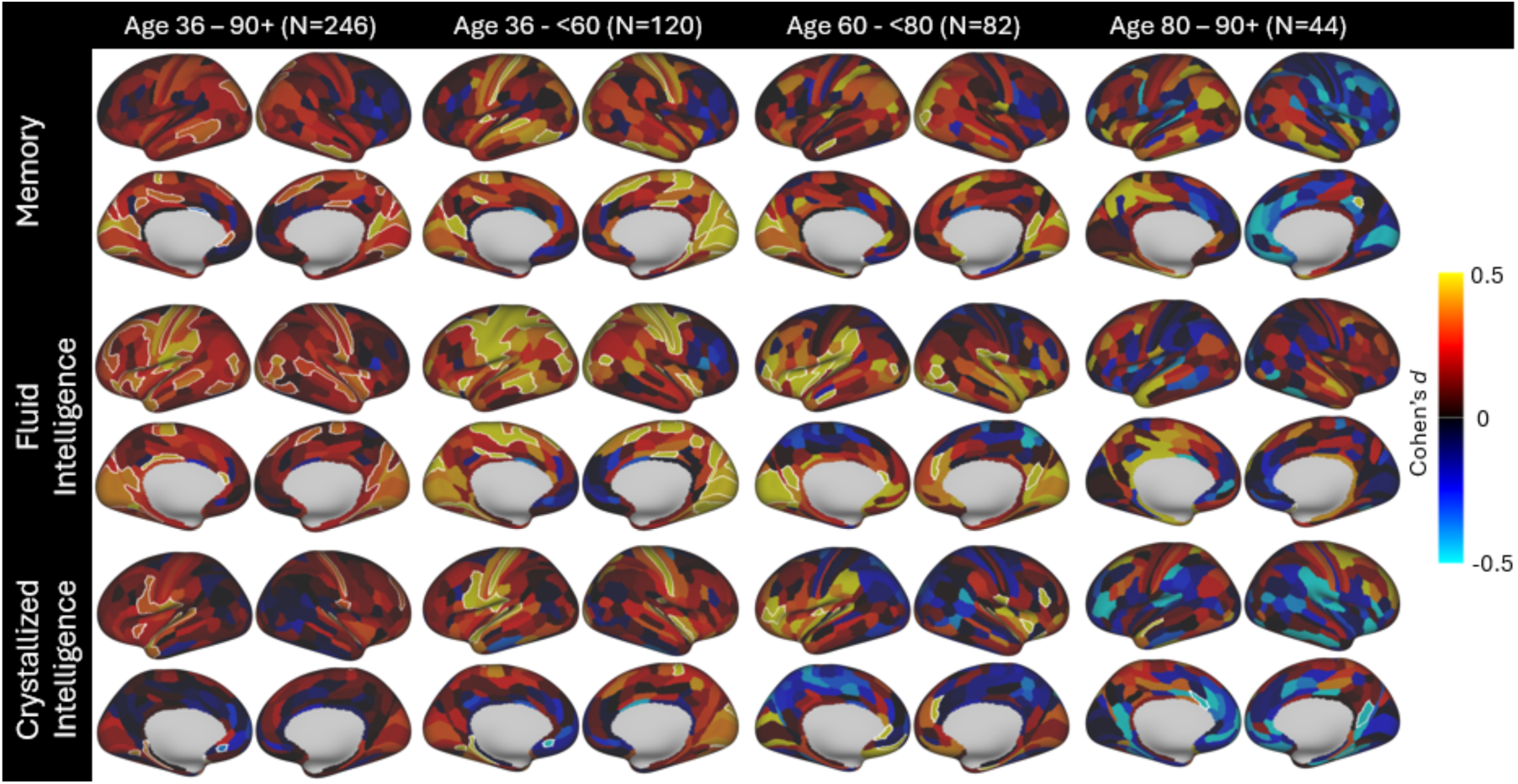
Spatial maps illustrate the group difference in cortical thickness residuals for each cortical area between top 10% and bottom 10% of cognitive performers, after adjusting for age, sex, and education. Effect sizes are expressed as Cohen’s *d* values and are signed according to the direction of group difference (warmer colors indicate cortical areas where cognitive high performers show greater cortical thickness; cooler colors indicate high performers with thinner cortex). Cortical areas surviving FDR correction (p < 0.05) are outlined in white.

**Figure 15.**
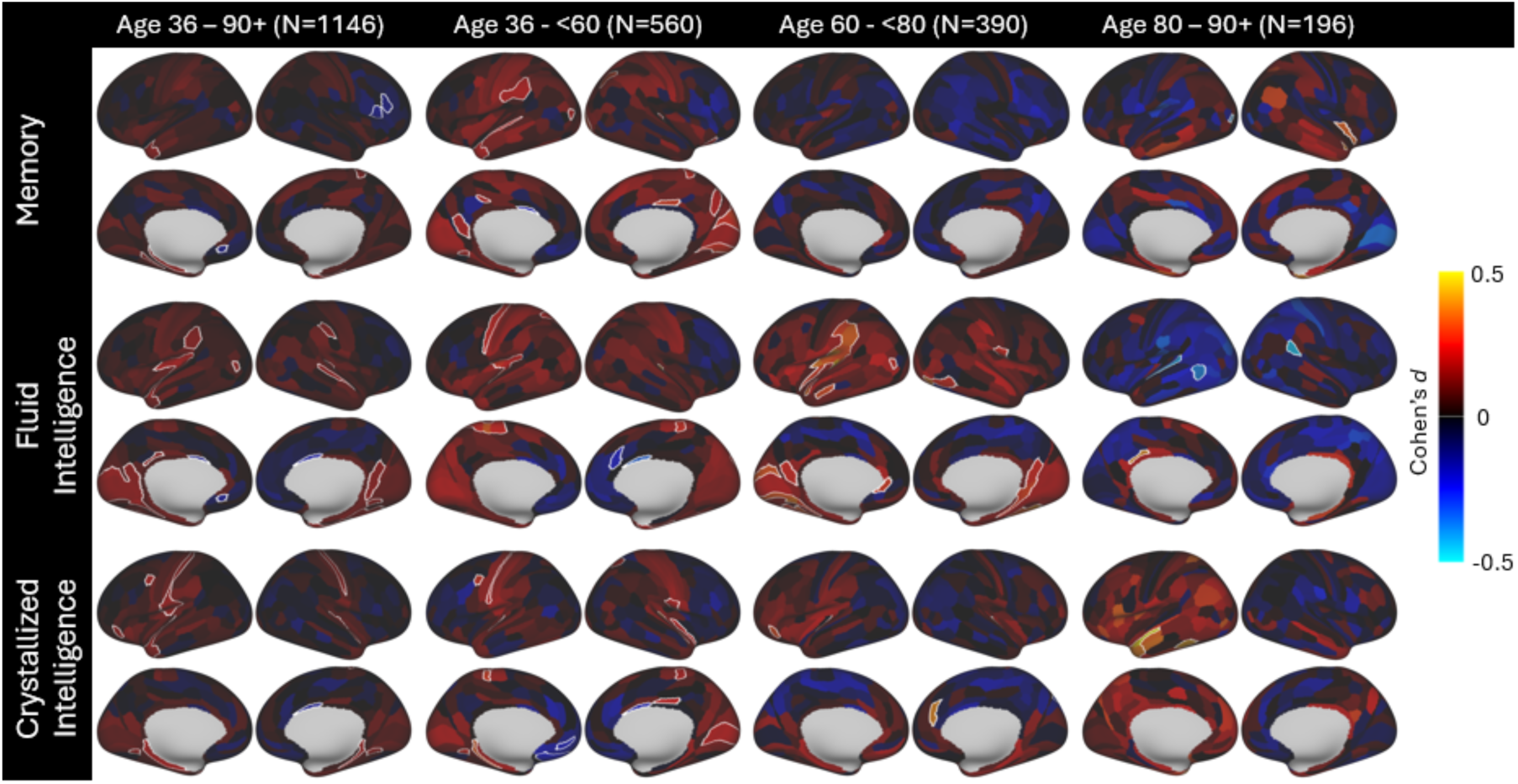
Spatial maps illustrate the group difference in cortical thickness residuals for each cortical area between top 50% and bottom 50% of cognitive performers, after adjusting for age, sex, and education. Effect sizes are expressed as Cohen’s *d* values and are signed according to the direction of group difference (warmer colors indicate cortical areas where cognitive high performers show greater cortical thickness; cooler colors indicate high performers with thinner cortex). Cortical areas surviving FDR correction (p < 0.05) are outlined in white.

Across the age groups, the middle-aged group (ages 36 - <60) showed strongest effects across all three cognitive domains involving cortical areas mostly in the somatomotor cortex, auditory cortex, visual cortex, and parietal regions. In the young-old group (ages 60–80), strongest cortical thickness differences were found between fluid intelligence high-/low-performers, particularly in the auditory cortex, inferior parietal regions, and visual cortex. Among the oldest group (ages 80–90+), cortical areas showing significant effects became more limited, even with the more extreme 10% performers.

Given the variability in cognitive test performance at a single time point, robustness tests were also conducted using longitudinally defined subsamples. The stable 25% subsample consisted of participants who consistently scored in the top or bottom 25% on both V1 and V2 assessments. Comparisons within this group revealed the most conservative, yet likely the most reliable, set of findings (Figure 16). Despite the significantly smaller sample size, several cortical areas that showed significant effects in the full baseline sample were replicated here.

**Figure 16.**
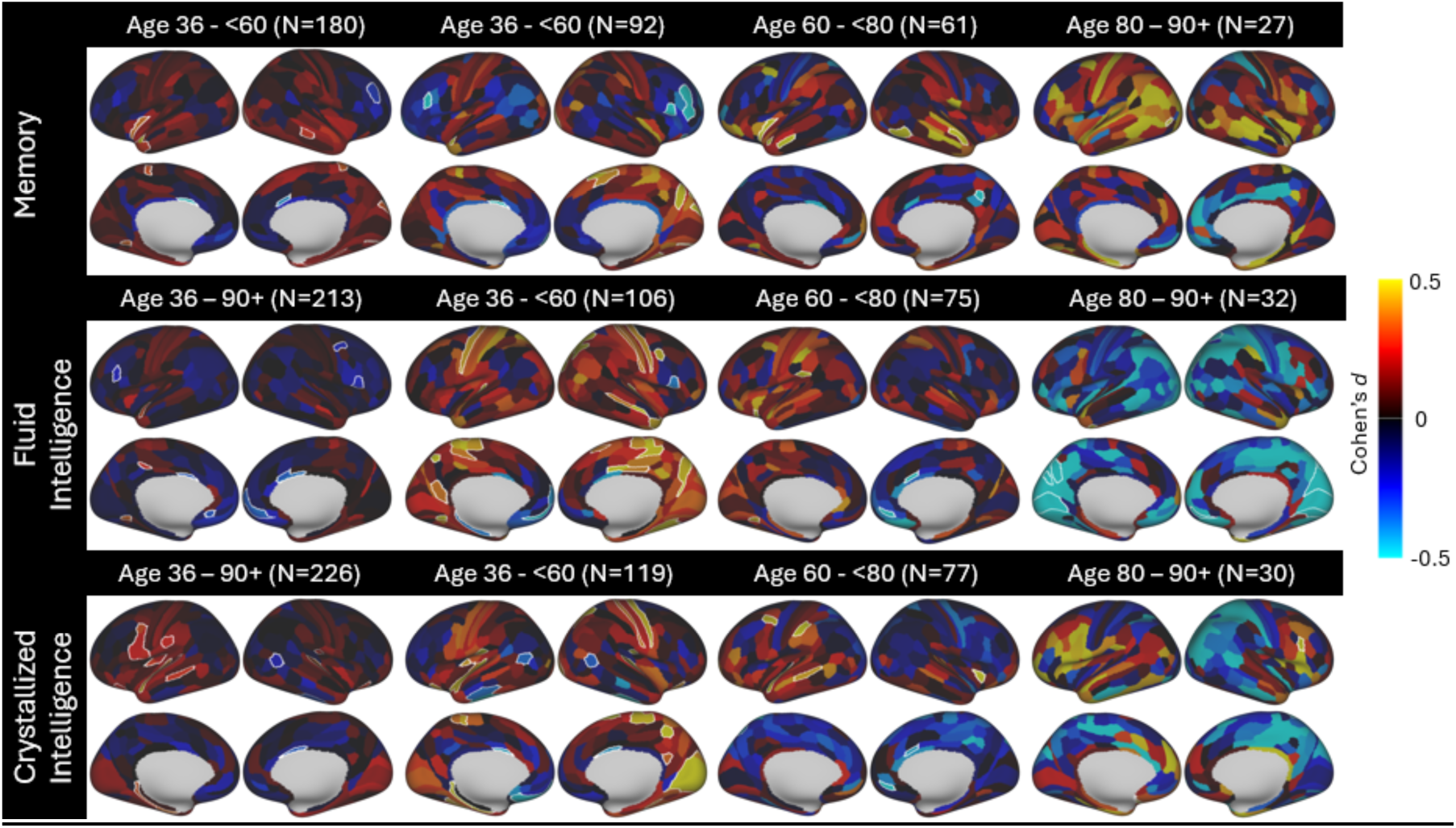
Spatial maps illustrate the group difference in cortical thickness residuals for each cortical area between stable top 25% and bottom 25% of cognitive performers defined using longitudinal data, after adjusting for age, sex, and education. Effect sizes are expressed as Cohen’s *d* values and are signed according to the direction of group difference (warmer colors indicate cortical areas where cognitive high performers show greater cortical thickness; cooler colors indicate high performers with thinner cortex). Cortical areas surviving FDR correction (p < 0.05) are outlined in white.

Specifically, middle-aged individuals with higher fluid intelligence scores showed greater cortical thickness in the somatomotor cortex (e.g., 4, 5L, 5m), visual cortices (e.g., VMV1, DVT, LIPv), as well as scattered cortical areas in the occipito-parietal cortex (e.g., 7Pm, IP1, POS1), posterior cingulate cortex (23d) and medial temporal lobe (PHA2). Similarly, individuals with higher crystallized intelligence exhibited greater thickness in somatomotor cortical areas (e.g., 1, 5m, OP1), auditory cortex (e.g., A1, MBelt, TA2), and primary visual areas (V1, V2, V3). Among participants in the 60–80 age group, the replicated findings were more limited and primarily involved auditory and language-related cortical areas (MBelt, STGa, TA2).

This smaller sample also yielded some counterintuitive findings where higher cognitive performers exhibited thinner cortex. A few of these effects were previously seen in the memory performance grouping using full baseline sample (i.e., frontal cortical areas of 33pr, 44, a10p, and IFSa in the 36–60 age group). Among the 80+ age group, such findings were observed primarily in the visual cortex, a pattern that was seen in the single-area analyses presented before (Figure 13).

Robustness tests were also conducted using an alternative longitudinally defined sample, which applied a different approach to leverage the longitudinal data. This method identified a subsample of baseline participants who were least likely to cross the regression line on repeated testing, thereby reducing classification instability over time. Although this approach does not provide the more cognitively distinct groups examined in our initial analyses (i.e., 25% and 10% groups), it provides a classification that can be considered robust against limited measurement reliability. This test slightly augmented the same patterns observed in the full 25% baseline sample comparisons (Figure 17).

**Figure 17.**
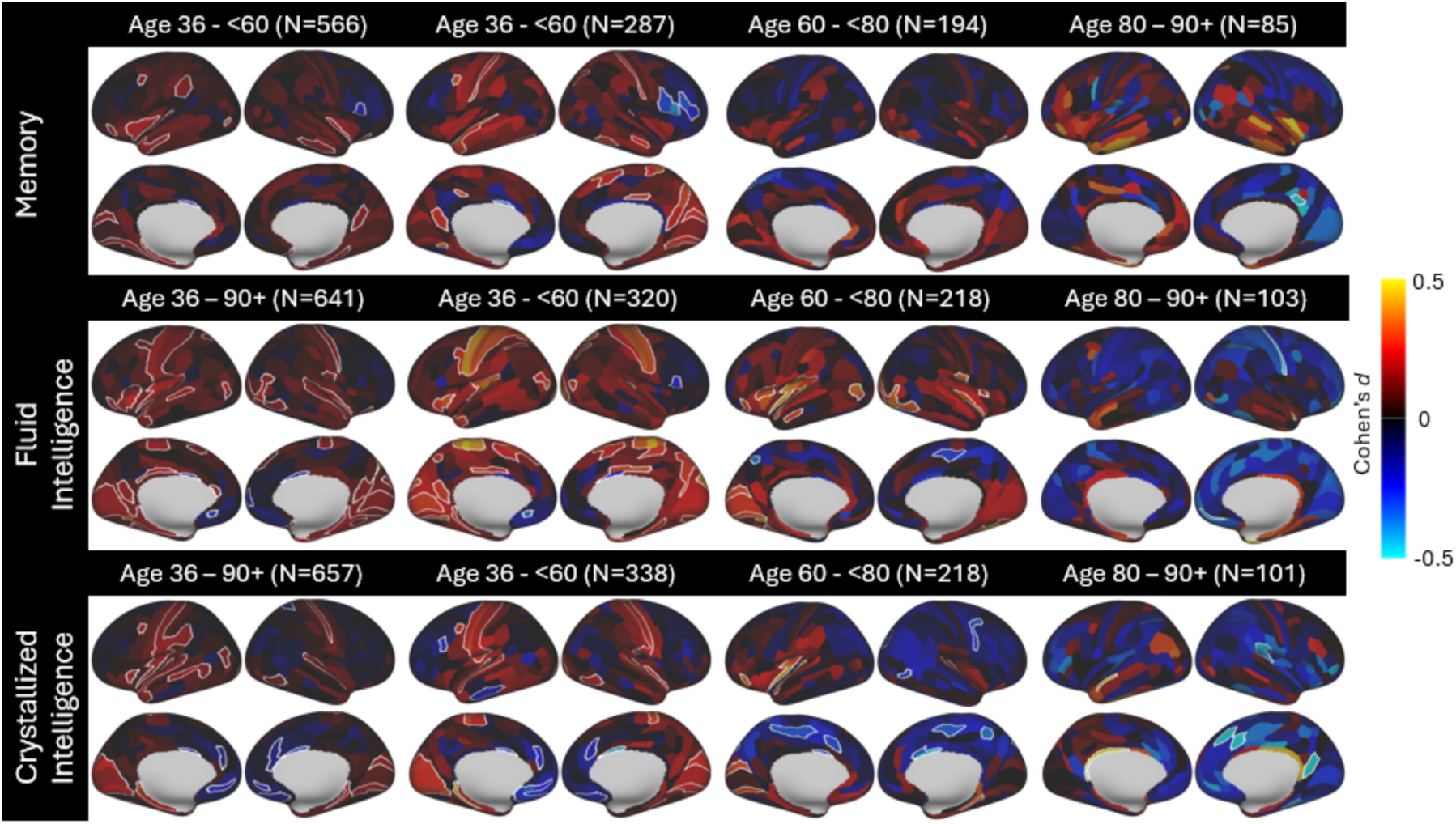
Spatial maps illustrate the group difference in cortical thickness residuals for each cortical area between stable top 50% and bottom 50% of cognitive performers defined using longitudinal data, after adjusting for age, sex, and education. Effect sizes are expressed as Cohen’s *d* values and are signed according to the direction of group difference (warmer colors indicate cortical areas where cognitive high performers show greater cortical thickness; cooler colors indicate high performers with thinner cortex). Cortical areas surviving FDR correction (p < 0.05) are outlined in white.

## 4. Discussion

The present study mapped whole-brain cortical thickness linked to cognitive performance in the large-scale HCP-A/AABC ‘typically’ aging cohort. By examining high- and low-cognitive performers within an otherwise cognitively healthy group, we aim to identify cortical areas of interest that may help define the mechanisms underlying superior cognitive performance in late life. Significant associations were observed between cortical thickness and cognitive factor scores representing memory, fluid intelligence, and crystallized intelligence, with the strongest effects found for fluid intelligence in younger age groups (36 - <60) and in female participants. These effects were most prominent in the somatomotor, visual and auditory cortices, though the strength of the association between cortical thickness and cognition generally decreased with advancing age. A series of secondary analyses generally supported the importance of a specific set of mostly unanticipated regions in these associations with cognition; however, idiosyncrasies in the results across analyses will require further evaluation to better understand those effects in the context of the primary study. For example, analyses by sex revealed stronger associations between cortical thickness and cognition in midlife females, particularly for fluid intelligence, while comparisons across the full age range revealed largely replicating spatial patterns between males and females. Thus, independent analyses suggest regional similarities in associations with cognition broadly, but effects do not necessarily replicate within a cognitive domain or age cohort.

Group differences in cortical thickness between ‘high’ and ‘low’ cognitive performers (percentile groupings) in this generally healthy cohort closely mirrored the results of the association analyses in the full sample. Specifically, the effects were strongest in both significance level and effect size among middle-aged adults (ages 36–59). In this age group, strong positive associations were observed between fluid and crystallized intelligence and cortical thickness in areas related to somatomotor, primary visual and auditory, and language functions. These findings appear robust, supported not only by their statistical significance and effect size, but also by their replication in multiple analytic subsamples. These patterns were also replicated in the younger-old age group (60–80), although with overall weaker effects. In the oldest group (80+), however, these effects became extremely limited. The most stringent robustness test based on longitudinally defined subsamples with stable performance classifications supported aspects of this pattern despite the notably smaller sample sizes. Contrary to expectations, these effects were disproportionately driven by the deviation from the typical population observed in the low-performing group, as opposed to being a unique feature of the high performers. However, a select set of cortical areas did exhibit such effects that may be associated with superior cognitive performance in our oldest group of the sample including the entorhinal cortex (EC) and cortical areas in posterior cingulate cortex (31pd) and superior temporal lobe (TA2).

The spatial pattern involving primarily motor and sensory regions were consistently observed across all three cognitive performance grouping thresholds (10%, 25%, and 50%), with the largest effect sizes seen in the 10% cutoff group, suggesting that group differences were driven by the most extreme performers.

The regional involvement observed across these comparisons resembles general aging-related patterns of cortical thinning in our sample (Figure 4). Importantly, our large HCP-A/AABC cohort enabled strict demographic matching of high- and low-performance groups, allowing us to minimize confounding by age and sex and to link group differences more confidently with cognitive performance. It also enabled separate analyses across three age bins (midlife through 90+). With this design, we observed a progressive attenuation of the cortical-thickness group effect, both in effect size and in the number of significant cortical areas, as age increased. This trend was further illustrated through single-area examinations of the most significant and highly replicated cortical areas, in which a convergence effect was observed. In these select cortical areas, differences in cortical thickness between performance groups diminished with age. In some cases, the direction of the effect even reversed with increasing age, with higher-performing individuals exhibiting thinner cortices.

Our results also implicated involvement of cingulate cortex (e.g., d32, 23d, d23b, 31pd) as highlighted by previous studies^11–13,15–17^, as well as scattered cortical areas in the MTL (e.g., EC, PeEc, PreS, PHA) that are vulnerable to Alzheimer’s disease^7,9^, though in a less consistent manner. However, these areas continued to emerge in the oldest subgroup and in the conservatively defined longitudinally-stable subgroup, where overall effects were the weakest. These areas may be related to the mechanism supporting superior cognitive performance in older adults that are different from the younger years, and warrant further exploration potentially through other imaging modalities of neural integrity.

Inferring from these cross-sectional data, this large lifespan cohort of cognitively healthy adults may suggest the role of premorbid differences in cortical brain structure (e.g., thicker cortex in younger high-performing individuals in regions sensitive to typical aging but less vulnerable to later life pathology such as Alzheimer’s disease pathology) in sustaining high cognitive performance.

Interestingly, a recent study of the APOE ε2 genotype (protective against AD) found larger gray-matter volumes in cognitively unimpaired carriers involving regions that overlap our findings^56^. In younger individuals (ages 36–59), the relationship between ’brain reserve’ (thicker regional cortex) and cognitive performance appears stronger and more widespread, particularly in primary and higher-order sensory processing. In contrast, this structure–performance relationship weakens with age, suggesting that cortical thickness may not reliably differentiate performance levels among older adults in this healthy cohort, or that older high-performers’ cognitive resilience may be supported by mechanisms different from that of the younger adults. In other words, it is possible that these cortical areas in the primarily sensory and motor regions reflect structural features of the brain driven by early-life determinants (e.g. genetics, prenatal and early developmental factors, education, SES, and environment)^57,58^ that contribute to the attainment of higher cognitive performance earlier in adulthood, but that these same features may not be sufficient to sustain superior performance into later life.

Indeed, our early analyses suggest that later life cognition may be more strongly linked to white matter integrity in this sample, as has been hinted at in previously reported studies of aging^15,59^. In addition, white matter integrity may also impact measurement of cortical thickness by influencing the measured distance between the gray/white matter borders defined by the image processing procedures. For example, regions showing thicker cortex in low performers show some overlap with regions prone to significant deterioration of juxtacortical white matter and alteration of the white matter intensities on T1 MRI^60^. These regions may result in the appearance of thicker cortex in individuals with white matter pathology, and thinner cortex when the white matter is preserved. Ongoing analyses will clarify the regional nature of white matter deterioration and the association between white matter integrity and the cognitive factors examined here. These results in combination may provide a new cortical morphometric pattern that differentiates those likely to have lower cognitive abilities in early life from those likely to perform at least typically and potentially show superior cognitive performance when other conditions are met.

A strength of the present study is the large normative community lifespan sample. Given this, we were able to examine a range of analyses (performance and age strata) which provided additional insights regarding the primary findings. The most basic finding, that a specific set of cortical areas are likely to be reduced in cortical thickness in low performing individuals in early life was largely supported across multiple secondary analyses and robustness tests, and also by the generally bilaterally symmetrical results (although formal laterality tests were not conducted). Separate models examining related constructs, such as linear models and group comparisons based on 10%, 25%, and 50% performance thresholds, revealed convergent effects in a similar set of cortical brain areas.

Additional analyses not reported but supporting the conclusions here included use of individual cognitive test scores in substitution for the factor scores to ensure that the factor process did not bias the results. Although some differences emerged, age- and sex-stratified analyses revealed considerable regional overlap, enhancing confidence in the replicability and robustness of the findings.

There are several limitations and considerations in this study. First, although a major strength of the HCP-A/AABC cohort is its large, lifespan-spanning sample, the number of participants in the oldest age range became more limited (Table 2), making comparisons across age bins less interpretable. Second, while the cognitive factor scores effectively summarize the primary dimensions of cognitive variance in the sample, they may not fully capture the complexity of cognitive functioning or the specific constructs most closely related to cortical brain structure. Similarly, the use of percentile-based groupings (e.g., 10%, 25%, and 50%) is useful for examining performance extremes but introduces somewhat arbitrary thresholds. These groupings may not optimally identify individuals with truly exceptional cognitive abilities. We are currently examining optimal methods for identifying individuals who deviate meaningfully from a normative distribution in terms of cognitive functioning.

Another key limitation is the cross-sectional nature of the study. Although we conducted robustness checks using longitudinally defined subsamples, cross-sectional designs limit the ability to reliably observe true age-related effects. Age-related changes in cortical thickness and cognition cannot be disentangled effectively from cohort effects or individual variability without longitudinal data.

Moreover, as with all behavioral measures, cognitive performance may be influenced by transient factors such as fatigue, motivation, or testing conditions. Relying on a single-timepoint assessment increases the risk of misclassifying individuals into high- or low-performance groups, which may reduce the sensitivity of the analyses and obscure true relationships. Future longitudinal research is essential to track intraindividual changes in both cognitive function and cortical brain structure over time and will allow distinction of developmental and premorbid effects from the effects of senescent aging. Ongoing work with the HCP-A/AABC cohort is incorporating longitudinal follow-up and novel cognitive constructs to better understand the dynamic interplay between brain morphology and cognitive aging, particularly in those exhibiting extreme cognitive preservation.

Additionally, several technical factors may impact the interpretation of any imaging results. As noted before, apparent ‘thickening’ of cortex with aging, or apparent thinner cortex in higher performers may partly reflect microstructural changes in adjacent white matter that shift the gray–white boundary used by surface pipelines. In this multi-site cohort, acquisition protocols and sequences were carefully standardized, and rigorous QC was applied; nonetheless, any residual site/scanner and image-quality variation may still impact data interpretation. Recently introduced approaches to quantifying macrostructural and microstructural properties simultaneously^10,61,62^ hold promise for more sensitive and specific structural indicators of cognitive aging and better handling of technical artifacts.

Finally, it should be noted that the histo/pathologic mechanisms underlying associations between cortical thickness and cognition are unknown and the specificity of these effects remain unclear. As a next step, we will incorporate biological and clinical covariates (e.g., APOE genotype, Aβ/tau biomarkers) to distinguish normative aging from disease-related processes.

The present findings provide a valuable foundation for understanding the complexity between brain morphology and cognitive aging, offering initial insights into how cognitive performance varies across the adult lifespan and identifying candidate regions and constructs for future studies focused on cognitive resilience and maintenance in late life.

## 5. Conclusion

We identified cortical areas where fluid intelligence, memory, and crystallized intelligence performance are associated with cortical thickness across the adult lifespan. Cognitive performance was linked to cortical thickness in an overlapping set of cortical brain areas, with top cognitive performers showing greater cortical thickness compared to bottom performers, particularly in midlife adults. The strongest associations were observed in somatomotor, visual, and auditory cortices during midlife (ages 36–59), while in older adults, associations weakened and became more variable. Effects were bilateral and largely replicated across different subsamples and analytic methods, supporting the robustness of these findings despite modest effect sizes. These results suggest that different mechanisms may underlie cognitive resilience across the adult lifespan. In midlife, higher cognitive performance appears to be supported by premorbid differences in cortical brain structure, particularly greater cortical thickness in primary and higher-order sensory and motor regions. In contrast, among older adults who are generally healthy, superior cognitive performance may be sustained by mechanisms not clearly reflected in cortical thickness measures.

