## Supplemental material for "Structural Brain Indicators of Cognitive Performance in Middle and Late Adulthood: The Human Connectome Project in Aging/Aging Adult Brain Connectome Cohort"

**Table S1. Factor analysis loadings using baseline neuropsychological data from 36-80-year-olds.**

| Neuropsychological Test | Memory | Fluid Intelligence | Crystallized Intelligence |
| --- | --- | --- | --- |
| RAVLT I | 0.58 |  |  |
| RAVLT II | 0.82 |  |  |
| RAVLT III | 0.89 |  |  |
| RAVLT IV | 0.88 |  |  |
| RAVLT V | 0.86 |  |  |
| RAVLT VI | 0.84 |  |  |
| MoCA Visuoconstructional |  | 0.34 |  |
| MoCA Naming |  |  |  |
| MoCA Digits |  |  |  |
| MoCA Letters |  |  |  |
| MoCA Subtraction |  |  |  |
| MoCA Sentence Repetition |  |  | 0.46 |
| MoCA Verbal Fluency |  |  |  |
| MoCA Abstraction |  |  |  |
| MoCA Delayed Recall | 0.40 |  |  |
| MoCA Orientation |  |  |  |
| NIH Toolbox Card Sorting |  | 0.71 |  |
| NIH Toolbox Flanker Task |  | 0.72 |  |
| NIH Toolbox List Sorting |  | 0.32 | 0.30 |
| NIH Toolbox Oral Reading |  |  | 0.72 |
| NIH Pattern Comparison |  | 0.68 |  |
| NIH Toolbox Picture Sequence Memory | 0.46 |  |  |
| NIH Toolbox Picture Vocabulary |  |  | 0.83 |
| Trail Making Test A (log-transformed) |  | 0.63 |  |
| Trail Making Test B (log-transformed) |  | 0.67 |  |

**Table S2. Additional demographic breakdown of the top and bottom performance groups for the full sample**.

| **Memory** | | | | | | | |
| --- | --- | --- | --- | --- | --- | --- | --- |
|  |  | **Top 25%**  **(n=283)** | **Bottom 25%**  **(n=280)** | **Top 50%**  **(n=580)** | **Bottom 50%**  **(n=566)** | **Top 10%**  **(n=124)** | **Bottom 10%**  **(n=121)** |
| Race | American Indian/Alaska Native  Asian  Black or African American  Hawaiian or Pacific Islander More than one race Unknown or not reported White | 0.0%  8.1%  4.6%  0.4%  3.9%  0.4%  82.7% | 0.4%  3.2%  26.4%  0.4%  5.0%  3.2%  61.4% | 0.2%  9.0%  7.9%  0.2%  3.6%  0.9%  78.3% | 0.4%  4.1%  22.6%  0.4%  4.2%  2.8%  65.5% | 0.0%  7.3%  4.0%  0.0%  3.2%  0.8%  84.7% | 0.8%  0.8%  30.6%  0.0%  5.8%  4.1%  57.9% |
| Ethnicity | Hispanic or Latino Not Hispanic or Latino  Unknown or not reported | 8.1%  91.9%  0.0% | 16.1%  83.9%  0.00% | 9.7%  90.2%  0.2% | 13.3%  86.6%  0.2% | 6.5%  93.5%  0.0% | 15.7%  84.3%  0.0% |
| APOE Genotype | APOE e4 +  APOE e4 –  APOE e2 +  APOE e2 –  Unknown | 20.8%  72.1%  10.2%  82.7%  7.1% | 21.8%  65.4%  8.2%  78.9%  12.9% | 20.3%  71.4%  11.7%  80.0%  8.3% | 20.1%  68.6%  11.0%  77.7%  11.3% | 20.2%  71.8%  8.1%  83.9%  8.1% | 18.2%  67.8%  7.4%  78.5%  14.0% |
| Education (years) | Mean (SD) | 17.98 (1.97) | 16.65 (2.51) | 17.92 (1.98) | 16.90 (2.43) | 18.13 (1.86) | 16.28 (2.78) |
| **Fluid Intelligence** | | | | | | | |
|  |  | **Top 25%**  **(n=283)** | **Bottom 25%**  **(n=280)** | **Top 50%**  **(n=579)** | **Bottom 50%**  **(n=567)** | **Top 10%**  **(n=124)** | **Bottom 10%**  **(n=122)** |
| Race | American Indian/Alaska Native  Asian  Black or African American  Hawaiian or Pacific Islander More than one race Unknown or not reported White | 0.0%  7.1%  4.2%  0.0%  4.2%  0.7%  83.7% | 0.7%  3.6%  32.9%  0.7%  5.7%  3.9%  52.5% | 0.2%  7.1%  5.7%  0.0%  3.8%  1.2%  82.0% | 0.4%  6.0%  24.9%  0.5%  4.1%  2.5%  61.7% | 0.0%  8.1%  4.0%  0.0%  3.2%  0.0%  84.7% | 0.0%  3.3%  39.3%  1.6%  5.7%  2.5%  47.5% |
| Ethnicity | Hispanic or Latino Not Hispanic or Latino  Unknown or not reported | 7.1%  92.9%  0.0% | 17.1%  82.5%  0.4% | 8.5%  91.4%  0.2% | 14.5%  85.4%  0.2% | 5.6%  94.4%  0.0% | 13.1%  86.1%  0.8% |
| APOE Genotype | APOE e4 +  APOE e4 –  APOE e2 +  APOE e2 –  Unknown | 20.5%  72.4%  12.7%  80.2%  7.1% | 22.9%  64.3%  10.0%  77.1%  12.9% | 19.0%  72.5%  12.3%  79.3%  8.5% | 21.5%  67.4%  10.4%  78.5%  11.1% | 20.2%  78.2%  14.5%  83.9%  1.6% | 23.8%  64.8%  9.8%  78.7%  11.5% |
| Education (years) | Mean (SD) | 17.99 (1.98) | 16.45 (2.63) | 17.91 (1.97) | 16.91 (2.44) | 18.04 (2.07) | 15.70 (2.85) |
| **Crystalized Intelligence** | | | | | | | |
|  |  | **Top 25%**  **(n=283)** | **Bottom 25%**  **(n=280)** | **Top 50%**  **(n=577)** | **Bottom 50%**  **(n=569)** | **Top 10%**  **(n=124)** | **Bottom 10%**  **(n=123)** |
| Race | American Indian/Alaska Native  Asian  Black or African American  Hawaiian or Pacific Islander More than one race Unknown or not reported White | 0.0%  5.3%  3.5%  0.0%  3.9%  0.0%  87.3% | 0.4%  8.6%  30.7%  0.7%  4.3%  4.6%  50.7% | 0.0%  4.9%  5.7%  0.0%  4.3%  0.7%  84.4% | 0.5%  8.3%  24.8%  0.5%  3.5%  3.0%  59.4% | 0.0%  4.0%  2.4%  0.0%  1.6%  0.0%  91.9% | 0.0%  8.1%  30.1%  0.8%  4.1%  7.3%  49.6% |
| Ethnicity | Hispanic or Latino Not Hispanic or Latino  Unknown or not reported | 4.2%  95.1%  0.0% | 20.4%  79.6%  0.00% | 6.4%  93.2%  0.3% | 16.5%  83.5%  0.0% | 2.4%  97.6%  0.0% | 26.8%  73.2%  0.0% |
| APOE Genotype | APOE e4+  APOE e4 –  APOE e2 +  APOE e2 –  Unknown | 19.1%  74.6%  12.4%  81.3%  6.4% | 23.6%  66.4%  11.4%  78.6%  10.0% | 20.3%  72.3%  11.4%  81.1%  7.5% | 20.2%  67.7%  11.2%  76.6%  12.1% | 21.8%  75.8%  8.1%  89.5%  2.4% | 20.3%  69.9%  12.2%  78.0%  9.8% |
| Education (years) | Mean (SD) | 18.50 (1.58) | 16.22 (2.67) | 18.04 (1.84) | 16.78 (2.48) | 18.95 (1.22) | 15.65 (2.89) |

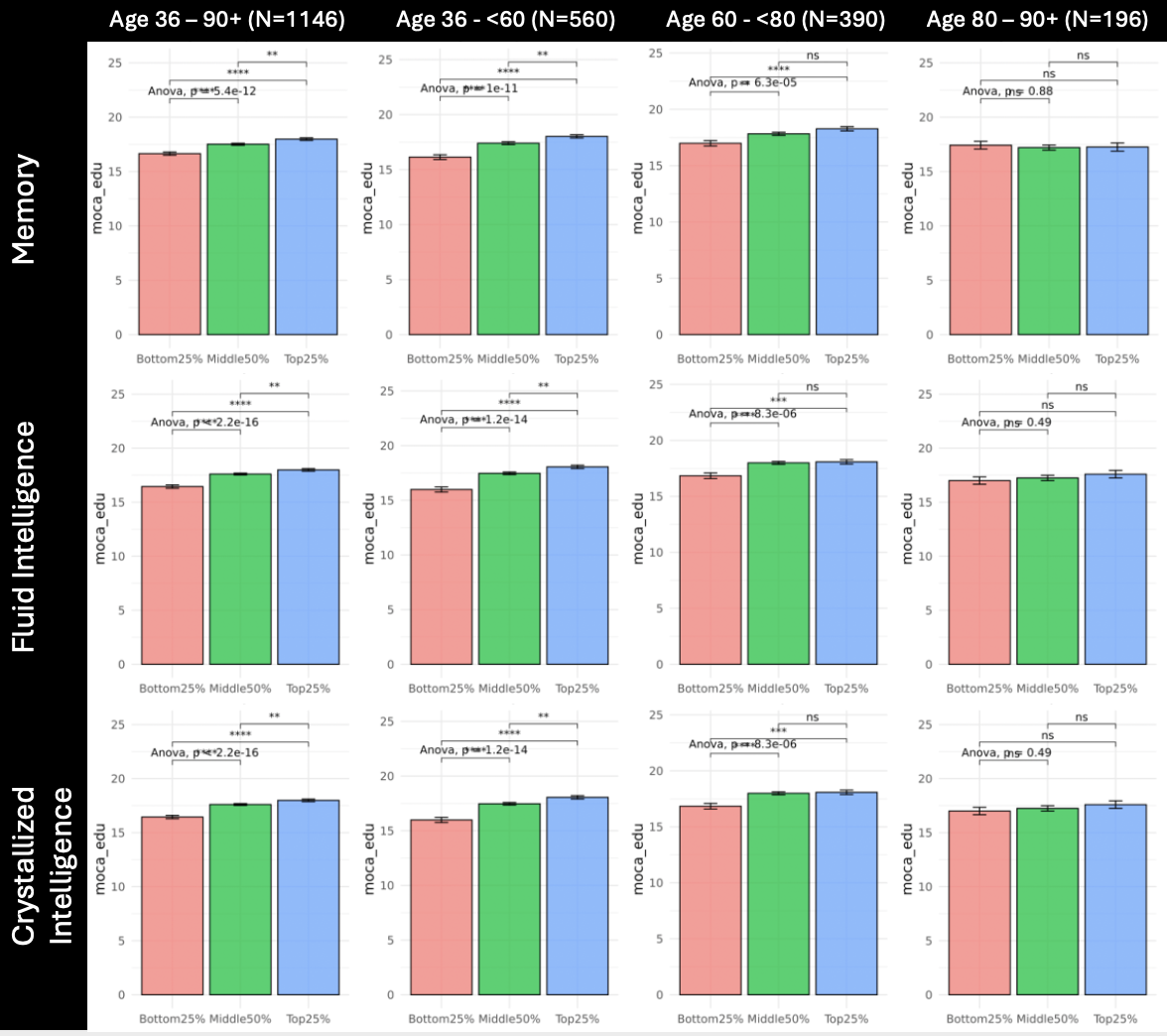
**Figure S1. Group differences in years of education between top 25% and bottom 25% of cognitive performers.**

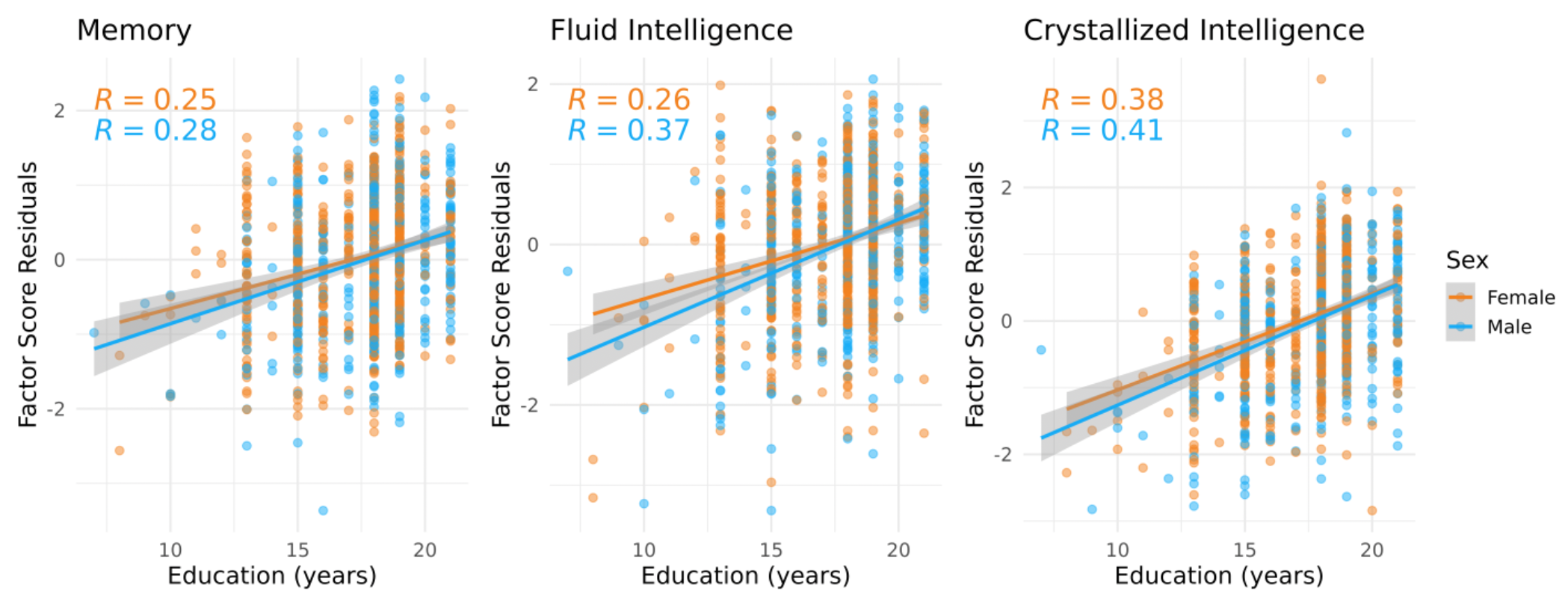

**Figure S2. Associations with education (year) for each factor score.**

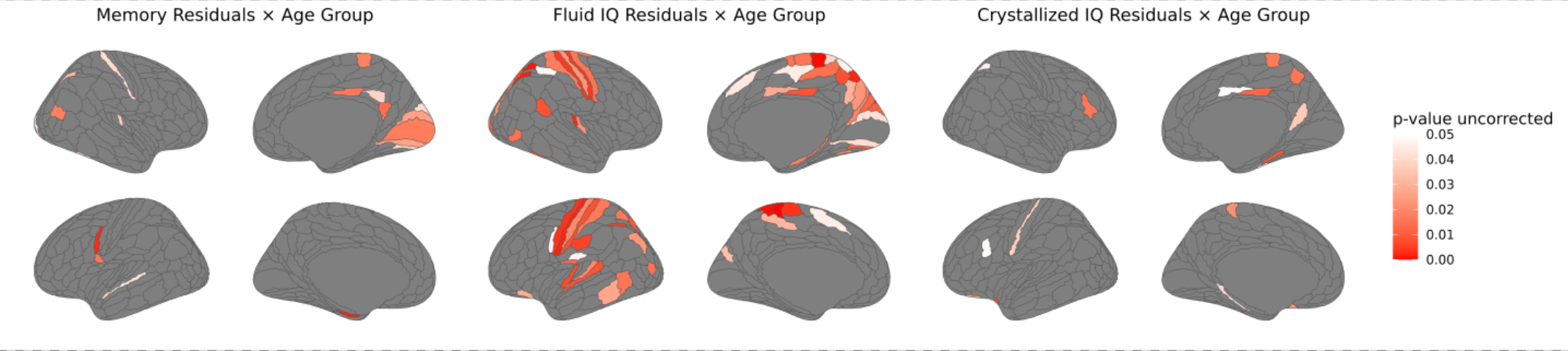

**Figure S3. Regions with significant interaction effects between cognitive performance and age group on cortical thickness.** The relationship between cognitive factor scores and cortical thickness changes significantly as a function of age group (i.e., 36-<60, 60-<80, 80+) in the colored regions.

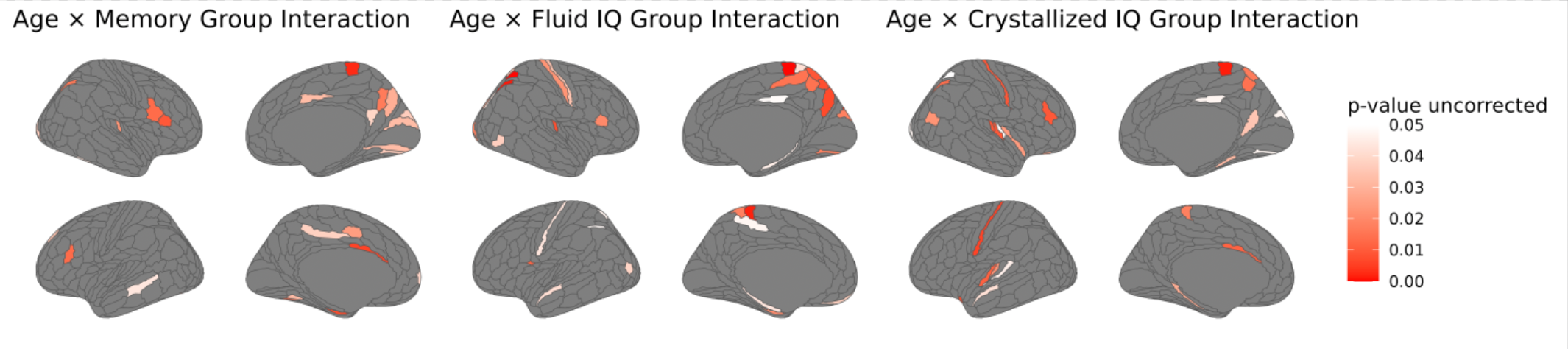

**Figure S4. Regions with significant interaction effects between age and cognitive performance group on cortical thickness.** The relationship between age and cortical thickness changes significantly as a function of cognitive performance group (i.e., top 25% vs. bottom 25%) in the colored regions.
